# Annotation-free Learning of Plankton for Classification and Anomaly Detection

**DOI:** 10.1101/856815

**Authors:** Vito P. Pastore, Thomas G. Zimmerman, Sujoy Biswas, Simone Bianco

**Affiliations:** Industrial and Applied Genomics, S2S - Science to Solution, IBM Research – Almaden, San Jose, CA USA; NSF Center for Cellular Construction, University of California San Francisco, San Francisco, CA USA

## Abstract

The acquisition of increasingly large plankton digital image datasets requires automatic methods of recognition and classification. As data size and collection speed increases, manual annotation and database representation are often bottlenecks for utilization of machine learning algorithms for taxonomic classification of plankton species in field studies. In this paper we present a novel set of algorithms to perform accurate detection and classification of plankton species with minimal supervision. Our algorithms approach the performance of existing supervised machine learning algorithms when tested on a plankton dataset generated from a custom-built lensless digital device. Similar results are obtained on a larger image dataset obtained from the Woods Hole Oceanographic Institution. Our algorithms are designed to provide a new way to monitor the environment with a class of rapid online intelligent detectors.

**Author Summary:** Plankton are at the bottom of the aquatic food chain and marine phytoplankton are estimated to be responsible for over 50% of all global primary production [1] and play a fundamental role in climate regulation. Thus, changes in plankton ecology may have a profound impact on global climate, as well as deep social and economic consequences. It seems therefore paramount to collect and analyze real time plankton data to understand the relationship between the health of plankton and the health of the environment they live in. In this paper, we present a novel set of algorithms to perform accurate detection and classification of plankton species with minimal supervision. The proposed pipeline is designed to provide a new way to monitor the environment with a class of rapid online intelligent detectors.

## Introduction

Plankton are a class of aquatic microorganisms, composed of both drifters and swimmers, which can vary significantly in size, morphology and behavior. The exact number of plankton species is not known, but an estimation of oceanic plankton puts the number between 3444 and 4375 [2]. Traditionally, plankton are surveyed using either satellite remote sensing, where leftover biomass is inferred indirectly through measurement of total chlorophyll concentration, or with large net tows via oceanic vessels [3], with subsequent microscopic analysis of the preserved samples. Satellite imaging methods are extremely accurate in terms of global geographic association and very useful for broad species characterization but may present practical challenges in terms of accuracy of the performed counts, species preservation and fine-grained characterization. The analysis of preserved samples, instead, allows for fine grained classification and accurate counting with narrow spatial sampling. More recently, real time observation of plankton species has been made possible by novel instruments for high-throughput *in situ* autonomous and semi-autonomous microscopy [4]. Such high-resolution imaging instruments make it possible to observe and study spatio-temporal changes in plankton morphology and behavior, which can be correlated with environmental perturbations. Sudden or unexpected changes in number, shape, aggregation patterns, population composition or collective behavior may be used to infer anomalous conditions related to potentially catastrophic events, either natural, like harmful algal blooms, or man-made, like industrial run offs or oil spills. Intelligent systems trained on curated data could help establish the characteristics of a healthy ecosystem and detect perturbations that may represent potential threats. More importantly, given the diversity of plankton morphology and behavior across species and the growing but still limited availability of high-quality labeled data sources, there is a need for algorithms which require minimal supervision to classify and monitor plankton species with a performance approaching that of supervised algorithms. Moreover, it is also desirable for such algorithms to aid the discovery of new plankton classes, which cannot generally happen with supervised classification techniques.

In this paper we propose a set of novel algorithms to reliably characterize and classify plankton data. Our method is based on an unsupervised approach to overcome the limits of supervised machine learning techniques, and designed to dynamically classify plankton from instruments that continuously acquire plankton images. First, we evaluate the performances of our algorithms on a mixture of ten freshwater plankton species imaged with a lensless microscope designed for in situ data collection [5]. Next, we evaluate the performance of our algorithms on an image dataset extracted from the Woods Hole Oceanographic Institution (WHOI) plankton database [6]. Machine learning methods are becoming a popular way to characterize and classify plankton [7]– [14]. A recent paper [15] explores the use of Convolutional Neural Networks to classify species of zooplankton, by introducing an architecture named ZooplanktoNet. The authors claim that their customized architecture can reach higher accuracy compared to standard deep learning configurations, like VGG, AlexNet, CaffeNet, and GoogleNet. In [16] and [17], the authors use an SVM based algorithm to classify species with high accuracy from the WHOI dataset. In a recent Kaggle competition contest (http://www.kaggle.com/c/datasciencebowl), the authors developed a deep learning architecture named DeepSea [18] to perform accurate classification of plankton collected with an underwater camera. In [19] the authors combine features obtained with multiple kernel learning to achieve higher accuracy than classic machine learning algorithms. However, all these advancements use supervised learning algorithms that rely on large labeled training sets which are very difficult and time consuming to create. Although recent computational advances may reduce the annotation burden for large biological datasets [20], a high-performance unsupervised learning algorithm can provide an alternative for real time unbiased in situ analysis.

## Results

### Plankton Classifier

We developed an unsupervised customized pipeline for plankton classification and anomaly detection, that we named *plankton classifier*. The pipeline, shown in Fig 1, is tested on a collection of videos containing ten fresh water species of plankton captured with a lensless microscope [5]. Each video is ten seconds long and contains one or more species. As the method is unsupervised, no labels are provided to the classifier during training. The plankton classifier consists of four modules: an **image processor, a feature extractor, an unsupervised partitioning module** and a **classification module**. The **image processor** examines each frame of video and generates cropped images of each plankter. The **feature extractor** examines each plankter image and generates a collection of features. The **unsupervised partitioning module** clusters samples by features into classes. The **classification** module comprises of a neural network-based **anomaly detector** to both perform classification based on the inferred labels and provide information to extend the database in an unsupervised manner. A sample is considered an anomaly with respect to a class if the extracted features are significantly different from the class average, as described below. The classification module also includes a standard neural network classifier, for performance comparison. See section materials and methods for a description of the modules in more details, along with the methods considered and tested that led to our final design.

**Fig 1.**
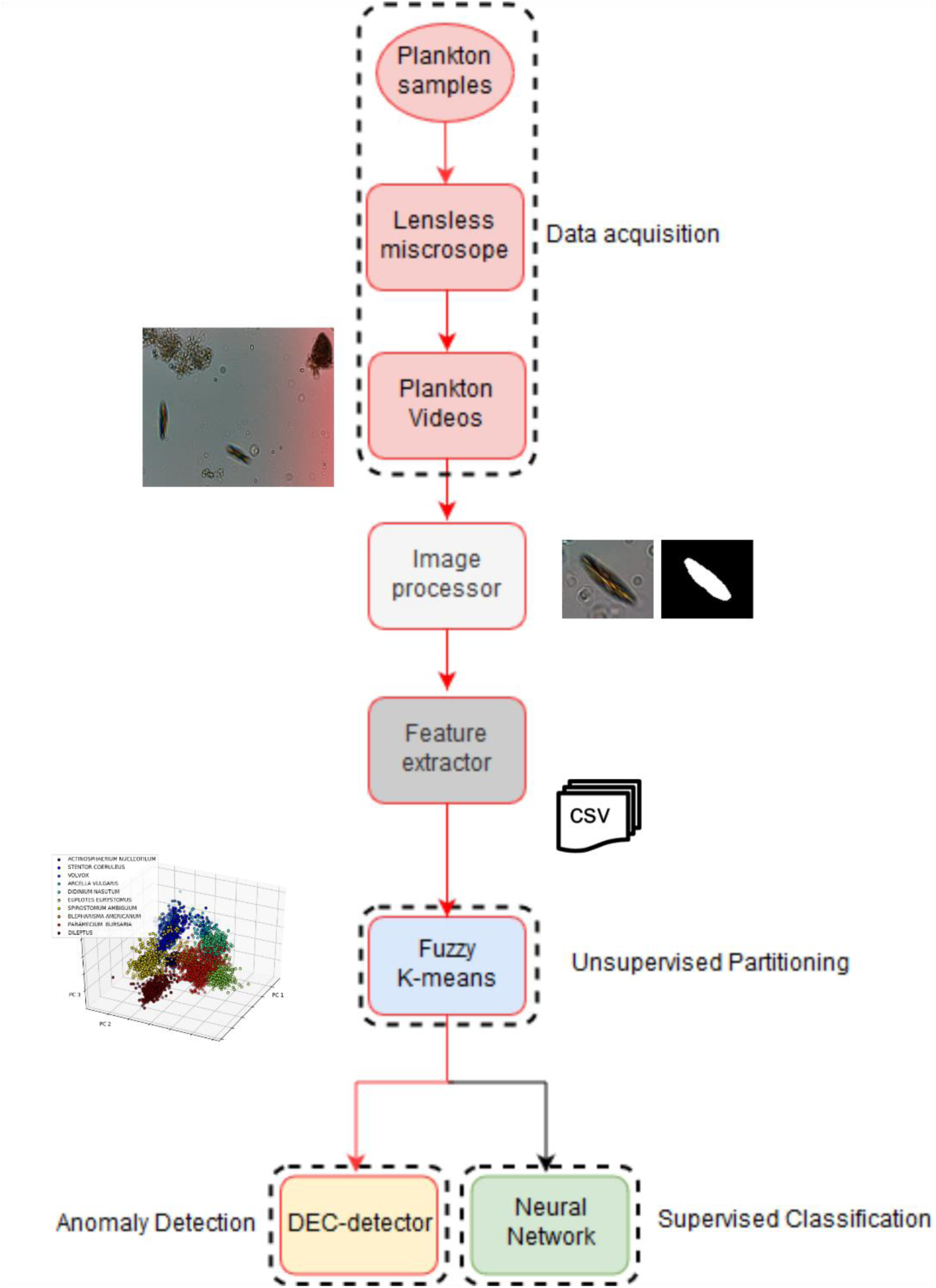
Schematic overview of the pipeline used to detect and classify plankton species with minimal supervision. Our preferred embodiment is represented by the red lines.

### Unsupervised partitioning performance

First, the plankton classifier examines each frame of an acquired video and generates cropped images of each plankter. A set of 131 features is then extracted, as described in Materials and Methods. The unsupervised partitioning module uses such features to place each plankton sample into one of Z classes. To automatically obtain the number of classes from the dataset, we have designed a custom algorithm based on partition entropy (see Materials and Methods). We evaluated the robustness of the implemented method on random subsets of the lensless dataset with different sizes, ranging from three to ten species. The box plot indicating the distribution for the estimated number of clusters Z among ten iterations can be observed in Fig 2e. The inferred number of classes, Z, is correctly identified in every case. A comparison of the performance of this algorithm against other existing methods is reported in the Supporting Information. Once we have obtained the number of clusters, we compared three clustering algorithms (see Supporting Information): k-Means, Fuzzy k-Means and Gaussian Mixture Model (GMM). Clustering accuracy is evaluated using purity (see materials and methods). The Fuzzy k-Means algorithm reaches a purity value of 0.934 (see Figs 2a, 2b), outperforming the standard k-Means (purity value = 0.887) and GMM [21] (purity value = 0.886). A posterior analysis of the results of the GMM reveals that this algorithm is not able to distinguish between *Blepharisma americanum* and *Paramecium bursaria*, due to their nearly identical appearance in the acquired videos. The Fuzzy k-Means algorithm is able to match the fuzziness exhibited by the plankton classes in parameter space which explains the lower accuracy of the crisp algorithms (k-Means and GMM). Therefore, we use the Fuzzy k-Means for our unsupervised classifier. A potentially important effect on the performance of any clustering algorithm is the class imbalance. The lensless microscope dataset is composed of 500 training samples for each of the ten considered species. To evaluate the impact of class imbalance, we performed the following experiment: We have built a dataset where the number of images of a species is a fraction (between 10% and 80%) of the number of images of the other species. We then evaluate the purity of this dataset and repeat the procedure for all the other species. Fig 2f reports the average performance over the ten datasets obtained as described above, as measured by the purity. The algorithm is always able to infer the correct number of species, without any overlap, with a minimum average purity value of 0.74 ± 0.09 (corresponding to 80% of class imbalance) and a maximum average purity value equal to 0.90 ± 0.08 (corresponding to 10% of class imbalance), with a maximum purity value of 0.972. This result shows that our pipeline can accurately cluster the data even in the case of strong class imbalance.

**Fig 2.**
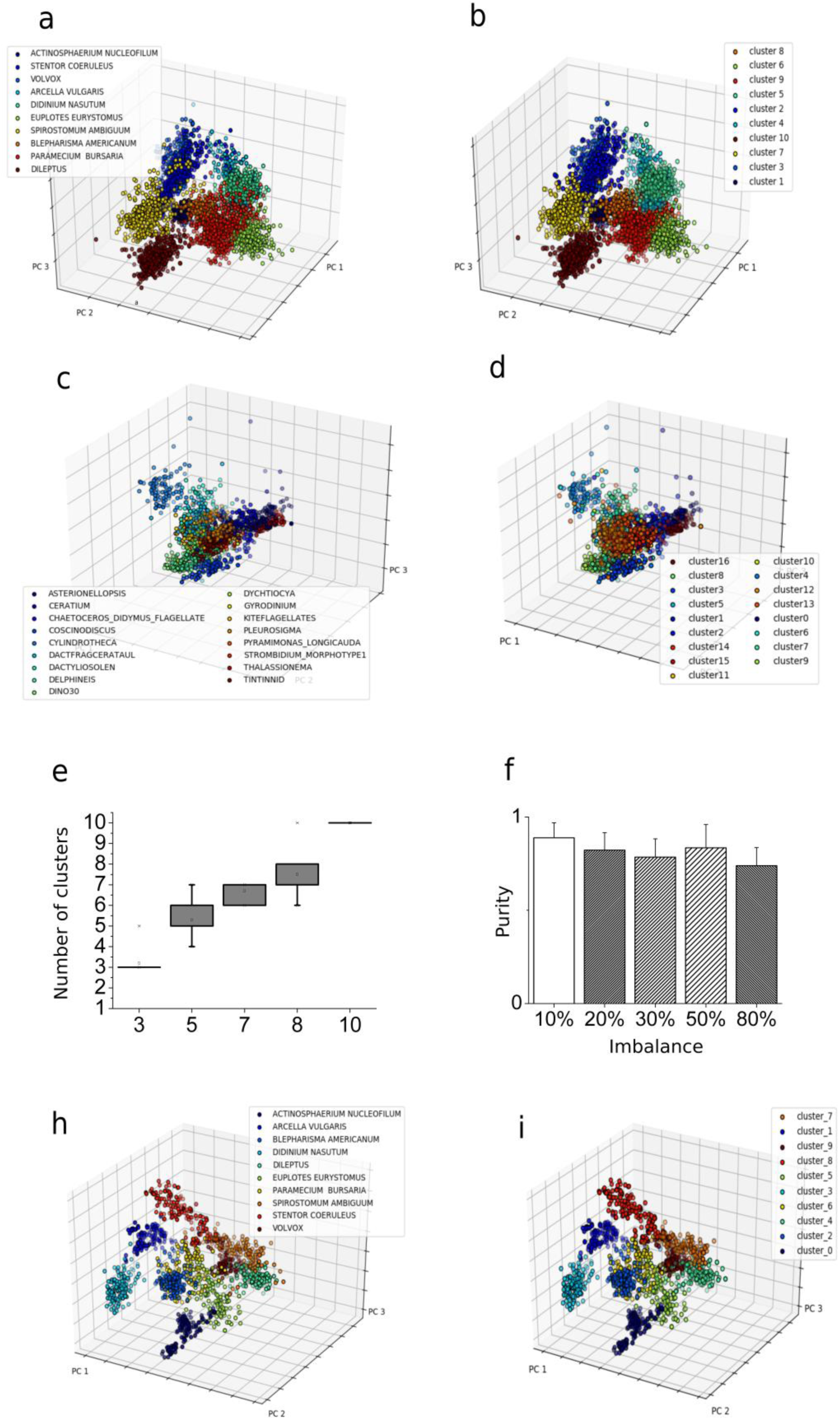
Unsupervised clustering results. **a**, **b** We performed a PCA analysis on the lensless digital microscope dataset to provide a graphical representation of the data distribution into the features space. We plot the first three principal components that account for ∼67% of the total variance. We assigned different colors to the different plankton species. **a** Species are assigned using ground truth labels. **b** Species are assigned to the most overlapping cluster resulting from the unsupervised partitioning procedure. **c, d** Same analysis and procedure applied on the WHOI dataset. **c** Species are assigned using ground truth labels. **d** Species are assigned to the most overlapping cluster, resulting from the unsupervised partitioning procedure. **e** Distribution of number of clusters computed using our PE algorithm for a random subset of species in the lensless microscope dataset. Results are reported for different initial number of species. **f** Effect of class imbalance. For each of the ten species included into the lensless microscope dataset, we simulated class imbalance by increasing the number of images available to the clustering algorithm for the considered species. **h, i** PCA analysis on the lensless digital microscope dataset provides a graphical representation of the data distribution into the deep features space. The unsupervised partitioning using deep features is highly accurate. The first three principal components are plotted and different colors to the different plankton species are assigned. **h** Species are assigned using ground truth labels**. i** Species are assigned to the most overlapping cluster resulting from the unsupervised partitioning.

### Algorithm performance on features extracted using deep feature extraction

Feature selection is an important part of any unsupervised learning pipeline. Indeed, hand engineering features introduces a degree of arbitrariness, which can be removed using a method of automated feature selection. Deep feature extraction, which consists in training a neural network architecture on either in- or out-of-domain data and use the last layer before prediction to extract features [9][22], is one such method. We trained the model described in section *Convolutional Neural Network (CNN) for deep features extraction* using the ten classes included in our lensless microscope dataset. The model reached 99% of training accuracy, 99% of validation accuracy and 98% of testing accuracy on the dataset obtained using our lensless microscope. Finally, the 128 neurons from the fully connected layers preceding the output are extracted and used as features for our pipeline. The PCA computed for the lensless microscope testing set among these features can be visualized in Fig 2h. Fig 2i shows the results of the unsupervised partitioning procedure. The underlying structure of the data set is very accurately captured, with a purity value of 0.98. Despite the fact that the accuracy obtained using deep feature extraction is slightly higher than the one obtained using the hand engineered features (purity of 0.980 vs 0.934), we decide to use the interpretable features described in Table 1. In fact, we think it is important that interpretability is maintained for the purpose of establishing a causal link between environmental perturbations and morphological modifications. However, for the purpose of organism classification, the customized deep feature extraction algorithm we implemented is a very viable alternative to the one proposed.

**Table 1:** List of morphological features extracted from the processed images. See Supporting Information for a detailed explanation.

| Class | Number | Description |
| --- | --- | --- |
| **Geometric feature** | 14 | Area (pixels), area (0-th order moment), perimeter, eccentricity, rectangularity, roundness, shape factor, width and height (minimum fitting rectangle), circularity, major and minor axis (fitting ellipse), equivalent diameter, convexity. |
| **Hu moments** | 7 | Hu moments computed from the normalized central image moments. |
| **Zernike moments** | 25 | Zernike moments up to order 5. |
| **Image intensity features** | 8 | Blue/green channels ratio, red/green channels ratio, red/blue channels ratio, gray levels histogram statistical features (skewness, kurtosis, mean value, standard deviation and entropy) |
| **Haralick features** | 13 | The first 13 features as proposed by Haralick in <sup>13</sup>, computed from the Gray Scale Co-occurrence Matrix (GSCM). |
| **Local binary patterns** | 54 | Local binary patterns summarize structures of the image comparing each pixel to its neighborhood |
| **Fourier descriptors** | 10 | Fourier descriptors are contour-based features invariant with respect to rotation, scaling and translation. |

### Classification

#### Supervised Classifier

At this stage of the pipeline, all samples have been assigned labels which have no correspondence to the actual plankton classes. We use the same trained clustering algorithm to classify the test samples, assigning each sample to the closest centroid. Using the trained Fuzzy k-means algorithm we reach a testing accuracy of 89%. Alternatively, one can use the labels obtained by our unsupervised partitioning algorithm to train a supervised classifier. We evaluated two algorithms: An Artificial Neural Network (ANN) and a Random Forest (RF) classifier. Our ANN architecture consists of a collection of classifiers, each trained to detect one plankton class. The RF approach consists in a set of decision trees to separate the training step samples into the correct classes.

For comparison, a simple ANN classifier is trained using the labels provided by the unsupervised partitioning algorithm. The ANN is a massive parallel combination of single processing units which can learn the structure of the data and store the knowledge in its connections [23]. See Materials and Methods for further information and for a detailed description of the implemented architecture. The network is very shallow, providing an efficient feature selection process. The ANN classifier reaches a validation accuracy of 99% and a testing accuracy of 94.5%. Figs 3c and 3d report the ROC curves and the confusion matrix obtained by testing the trained ANN classifier on our ten species plankton dataset. The ROC curves are close to a perfect classifier and the confusion matrix is almost diagonal with minor overlap between two pairs of species: *Blepharisma americanuum-Paramecium bursaria* and *Spirostomum ambiguum*-*Stentor coerouleus*. This misclassification is primarily due to the similarity in the shape, size and texture of the two pairs of species, influencing both the unsupervised training clustering and the subsequent testing of the supervised classifier.

**Fig 3.**
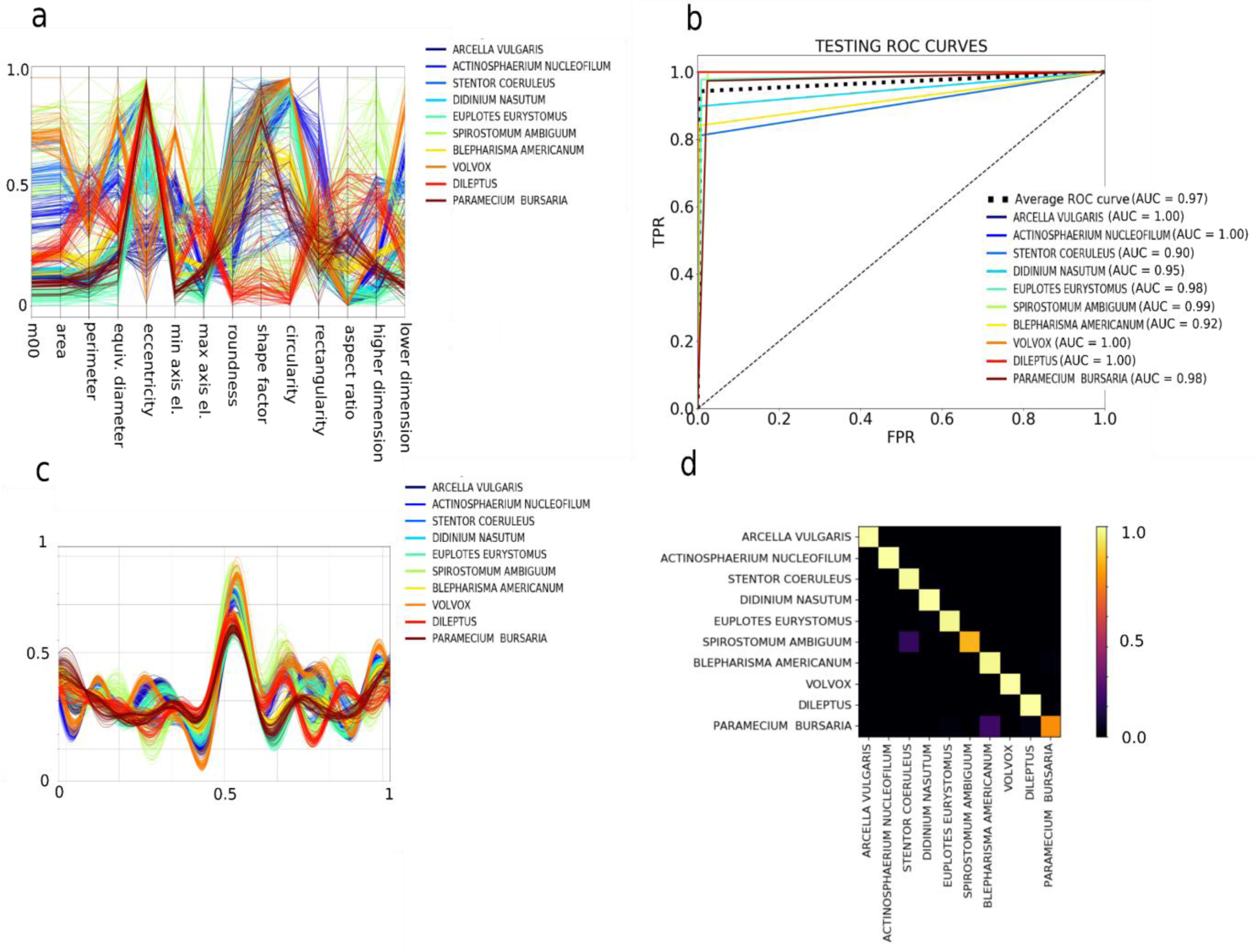
Feature space representation and classification performances. **a, b** Multidimensional visualization of the geometric subset of the ten species in the lensless microscope dataset, obtained using the following methods (see Supporting Information): **a** Andrew’s curve. **b** Parallel coordinates. **c** ROC curves obtained for the neural network classifier trained on the labels provided by the clustering algorithm for the lensless microscope dataset. **d** Corresponding confusion matrix.

An alternative classifier method employs a Random Forest (RF) approach, a popular ensemble learning method used for classification and regression tasks.

We train an RF algorithm using the labels provided by the unsupervised classifier and reach an accuracy of 94%. For comparison, we train the same RF algorithm using the actual labels (ground truth) of the training set and reach an accuracy around 98%, proving that our unsupervised classification approach performs comparably well with respect to the correspondent supervised approaches for the trained classifier. Since the ANN performs marginally better than the RF classifier, we propose the former for a pipeline. In the next section, we will present an alternative classification method

### Anomaly Detector

When deployed in the field, microscopes will encounter species that have never been seen before, so it is essential that such samples are detected and correctly identified as anomalies. For a given class, a sample is considered an anomaly if the sample features are significantly different from the feature average for the class. Algorithms for anomaly detection based on the separation of the features space have been successfully used to identify the intrusion in computer networks for security purposes [24]. Two anomaly detectors are implemented and compared; a state of the art one-class SVM^15^ and a customized neural network we call a Delta-Enhanced Class (DEC) detector that combines classification with anomaly detection. The one-class SVM algorithm uses a kernel to project the data onto a multidimensional space and can be interpreted as a two class SVM assigning the origin to one class and the rest of the data to another class. It then solves an optimization problem determining a hyperplane with maximum geometric margin, i.e., a surface where the separation between the two sets of points is maximal, that will be used as decision rule during the testing step.

A customized one-class SVM is implemented by normalizing the testing samples using the training data belonging to a single class. In this way, there will be a significant difference in the absolute value obtained for the anomaly (out-of-class) samples compared to the in-class samples, improving the accuracy of the SVM. The one-class SVM so designed reaches an average testing accuracy of (93.5 ± 6.0) %, with high accuracy in both anomaly detection and classification.

We now describe an alternative ANN-based approach that simultaneously performs classification and anomaly detection. As demonstrated above, a single layer ANN is able to satisfactorily classify plankton data from our in-house dataset. However, to effectively approach the anomaly detection step, we designed a deep neural network called Delta-Enhanced Class (DEC) detector (see materials and methods for further details). One DEC detector must be trained for each of the training species. Therefore, we train ten DEC detectors, one for each of the species of plankton identified in the unsupervised learning step. This procedure affords excellent accuracy on both classification and anomaly detection, on both real and simulated plankton data (see Fig 4), with an average testing accuracy on real data of 98.8 ± 2.4 %, an average anomaly detection testing accuracy of 99.2 ± 0.7 % and an average overall testing accuracy of 99.1 ± 0.9 % (see Fig 4b for details). The confusion matrices in Fig 4a demonstrate the discrimination power of our algorithm. The DEC detector outperforms the alternative one-class SVM classifier in both supervised (average accuracy equal to 95%) and unsupervised (average accuracy equal to 93.5%) configurations. It is worth reporting that the unsupervised one-class SVM reached a minimum overall accuracy of 79%, compared to 97.2% for the DEC detector (minimum values correspond to *Paramecium bursaria* detector). To test the overall performance of our method, we produce a dataset of surrogate plankton organisms. For each different species, we test the corresponding DEC detector architecture using a surrogate species created with a feature-by-feature weighted average of all the species in our dataset. Starting with a uniform weight distribution, we increase the weight for the species corresponding to the trained DEC detector architecture up to 0.9 (steps of 0.1), obtaining 9 different surrogate species (see Fig 4d for an average parallel coordinates plot, showing the resulting distributions for the species *Spirostomum ambiguum*). The aim of this robustness test is to simulate the acquisition of an unknown species, whose features are increasingly closer to the features of the class correspondent to the detector, up to a maximum of 90% similarity. As Fig 4e shows, our classifier can recognize the synthetic species as an anomaly with an average accuracy higher than 98% if the similarity between the synthetic and the real species is up to 30%, and it can maintain an average accuracy of over 82.6% if the species similarity is up to 50%. Accuracy of anomaly detection severely decreases if the species similarity is over 50%, reaching the minimum value of 37.5%.

**Fig 4.**
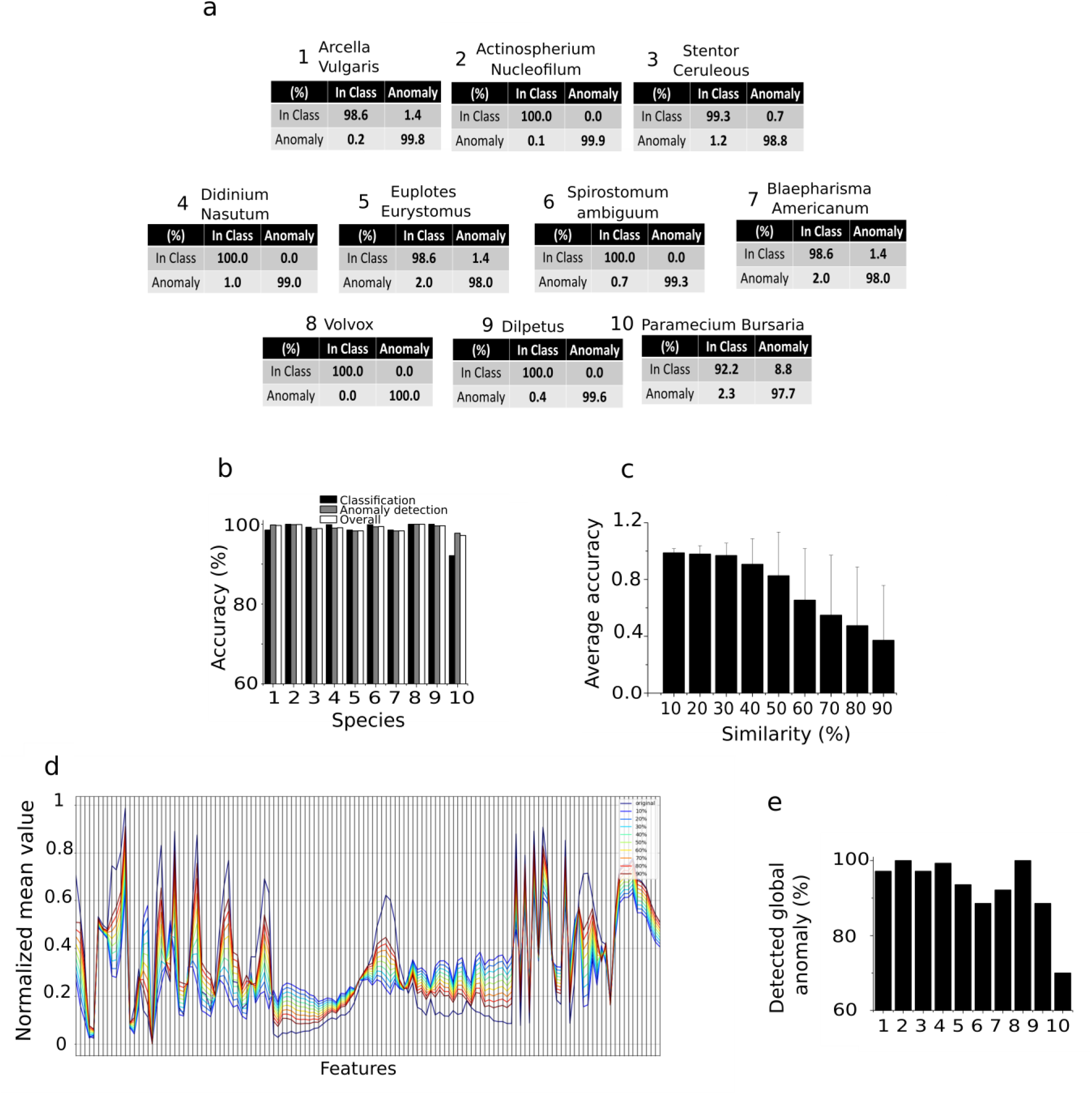
Delta-Enhanced Class detector performances and results. **a** Confusion matrix corresponding to each of the ten neural networks trained on the lensless microscope dataset. **b** Overall testing accuracy performances for each of the ten testing classes. The number used on x axis to label each species correspond to the species number in panel **a**. **c-d** DEC detector anomaly detection performances tested on in silico generated data. **d** Testing accuracy performances for varying percentage values of in silico species similarity with the trained species. **e** Example of average features space parallel coordinates plot for the in-silico species obtained using the species *Spirostomum Ambiguum*. By increasing the similarity, the features of the surrogate species approach the features of the real species, resulting in an increased average anomaly misclassification rate, decreasing the overall accuracy levels. **e** Detection of unknown species. The panel shows the percentage of samples detected by all the DEC detectors as anomaly, when removing one training species from the set, for each of the ten training species. These numbers reflect the level of accuracy of the proposed algorithm in detecting unseen species. The number used on x axis to label each species correspond to the species number in panel **a**.

### Plankton classifier performance on the WHOI dataset

The WHOI provides a public dataset comprising millions of still monochromatic images of microscopic marine plankton, captured with an optical Imaging FlowCytobot (https://mclanelabs.com/imaging-flowcytobot/). To use this dataset as a benchmark to test our unsupervised classifier, we extract a set of 128 features from a collection of 40 species of plankton (100 images per species, randomly selected), using both the segmented binary image and the portion of the gray-scale image containing the plankton cell body. A full description of the species selection process is reported in the Supporting Information. The features set is identical to the one used for the lensless microscope dataset, except for the absence of three-color features, as the lensless microscope is a color-based sensor, while the Imaging FlowCytobot is monochromatic. Figs 2c, 2d show the results of our pipeline applied on the normalized features set. The algorithm reaches an overall purity value of 0.715 for the 40 WHOI species that we selected. The ability of our pipeline to distinguish between inter-species plankton morphology can be further observed comparing Fig 2c, which represents the PCA space corresponding to a subset of 18 of the 40 species for the ground truth dataset, and Fig 2d, which represents the corresponding PCA space resulting from the unsupervised partitioning algorithm. A complete PCA representation for the 40 species can be found in Supporting Information. We trained a random forest algorithm using the labels provided by the unsupervised partitioning with a train-test ratio of 80:20, obtaining a classification accuracy around 63%. For comparison, we have trained a supervised random forest algorithm using the ground truth labels on the extracted features, obtaining a classification accuracy around 79%.

### The plankton classifier can reveal unseen species

We have demonstrated that our DEC neural networks are able to classify a sample as either a training class (i.e., the plankton species used to train the detector) or as an anomaly. If a sample is discarded by all the implemented detectors, it could either represent an intra-species anomaly (i.e., species included into the training set) or a sample belonging to an unseen species (i.e., species not included in the training set). The former represents the basis for using the proposed pipeline for real-time environmental monitoring, and its implications are discussed in the next section. We now test the potential of our pipeline to detect new species. We remove one class from our unsupervised partitioning ensemble set, consider it as never before seen and compute the number of testing samples detected as anomaly by all the remaining DEC detectors. This number indicates the algorithm accuracy in detecting new species. We repeat the procedure for each class. The average detection accuracy is 98.3 ± 10.1 % (see Fig 4e), demonstrating the ability of the pipeline to detect the presence of a new species. If two or more unseen species are detected, they will be stored as anomalies. As this group of anomalies grows, a human expert may determine offline the actual labels for these new species, thus allowing a DEC detector to be trained for each new species. Alternatively, the samples corresponding to unseen species may be clustered and classified by the unsupervised partitioning step of our pipeline, reducing the number of new species that must be examined by a human.

## Discussion

The plankton classifier described in this paper provides the foundation for a robust, accurate and scalable mean to autonomously survey plankton in the field. We have identified interpretable and non-interpretable image features that work with our algorithms to perform an efficient clustering and classification on plankton data using minimal supervision and with a performance accuracy comparable to supervised learning algorithms [16]. Instead of labeling thousands of samples, an expert need only identifying one member of cluster to label all the samples of the cluster.

We introduced a neural network that performs classification by learning the shape of the feature space and uses this information to identify anomalies. The network uses a novel unbiased methodology of feature-to-feature comparison of a test sample to a random set of training samples. While most of the existing classification methods require various degrees of user input, our method is automated, without sacrificing performance accuracy or efficiency.

All features the plankton classifier relies upon are extracted from static images. However, our custom lensless microscope captures 2D and 3D dynamic of plankton. While this dynamic information is not considered in the analysis presented here, motion data can increase the dimensionality of the feature space, by adding spatio-temporal “behavioral” components, and may improve the performance of classifiers and anomaly detectors. This is particularly valuable in cases where species have considerable overlap in morphology feature space, as seen with *Blepharisma americanuum* and *Paramecium bursaria*, and *Spirostomum ambiguum* and *Stentor coerouleus*, shown in the confusion matrices in Fig 3d. Currently, existing large plankton datasets, like the WHOI used in our validation experiments, are based on static images, but as the cost of video-based *in situ* microscopes drops and their deployment increases, we believe datasets that include spatio-temporal data will become available and the use of such features will gain importance.

Deploying smart microscopes capable of real-time continuous monitoring will give biologist an unprecedented view of plankton *in situ*. The adoption of an unsupervised unbiased pipeline is a significant step ahead in the development of a real-time “smart” detector for environmental monitoring. Several high-resolution acquisition systems for real-time plankton imaging already exist [25] and could adopt the pipeline proposed into this paper. Fig 5 shows a high-level representation of a continuous environmental monitoring system in the form of a flow chart, showing an example of how the detector could be coupled to the computational pipeline we designed. Once the descriptors have been extracted from the acquired videos, it is possible to use them to build a set of DEC detectors. It is important to stress that the size of the data likely to be acquired, or already present in the databases, makes neural networks the obvious choice to carry out the analysis due to their unsurpassed scalability. Our newly designed and customized DEC detector neural architecture for plankton classification and anomaly detection is a functional and efficient example of such algorithm. Moreover, neural algorithms can infer non-linear relationships between features (input) and correlate them with the class description (output) without making any assumptions on the underlying learning model. Hence, the classification depends only on the extracted features. Every time the network identifies a species belonging to a specific class, the average set of morphological features is then updated, thereby further qualifying the class morphology phase space. If an anomaly is detected, it may be sent to an expert for a supervised examination. The expert will determine whether that sample could be a species not represented in the training set, or if it belongs to an existing training class, but its morphological features deviate significantly from the average features space of the corresponding class. In the former case, a new smart detector will be trained offline, so that the training set is dynamically expanded, and the system will provide a continuous monitoring of the aquatic environment using the human expert-in-the-loop paradigm. In the latter case, the identified anomalies may represent local environmental perturbations, either natural or man-made. Further work is needed to assess the validity of such hypothesis. An additional re-training step may be necessary to update the algorithms. Our pipeline is based on local analysis using a low powered device, capable of image capture and processing, classification and anomaly detection. Coupling such platform with a local (laptop, server) or cloud-based system where the training step may occur could provide the flexibility and resources needed to close the loop and generate the training data the low power platform can use for classification. Examples of systems that use this paradigm are already present in the literature [26], and we hope the availability of computational paradigms like the one we propose may increase the research in the field. A high-resolution plankton acquisition system placed in the water and powered with our unsupervised pipeline may enable the development of real time continuous smart environmental monitoring systems that are fundamentally needed to stakeholders and decision-making bodies to monitor plankton microorganisms and, consequently, the entire aquatic ecosystem [27].

**Fig 5.**
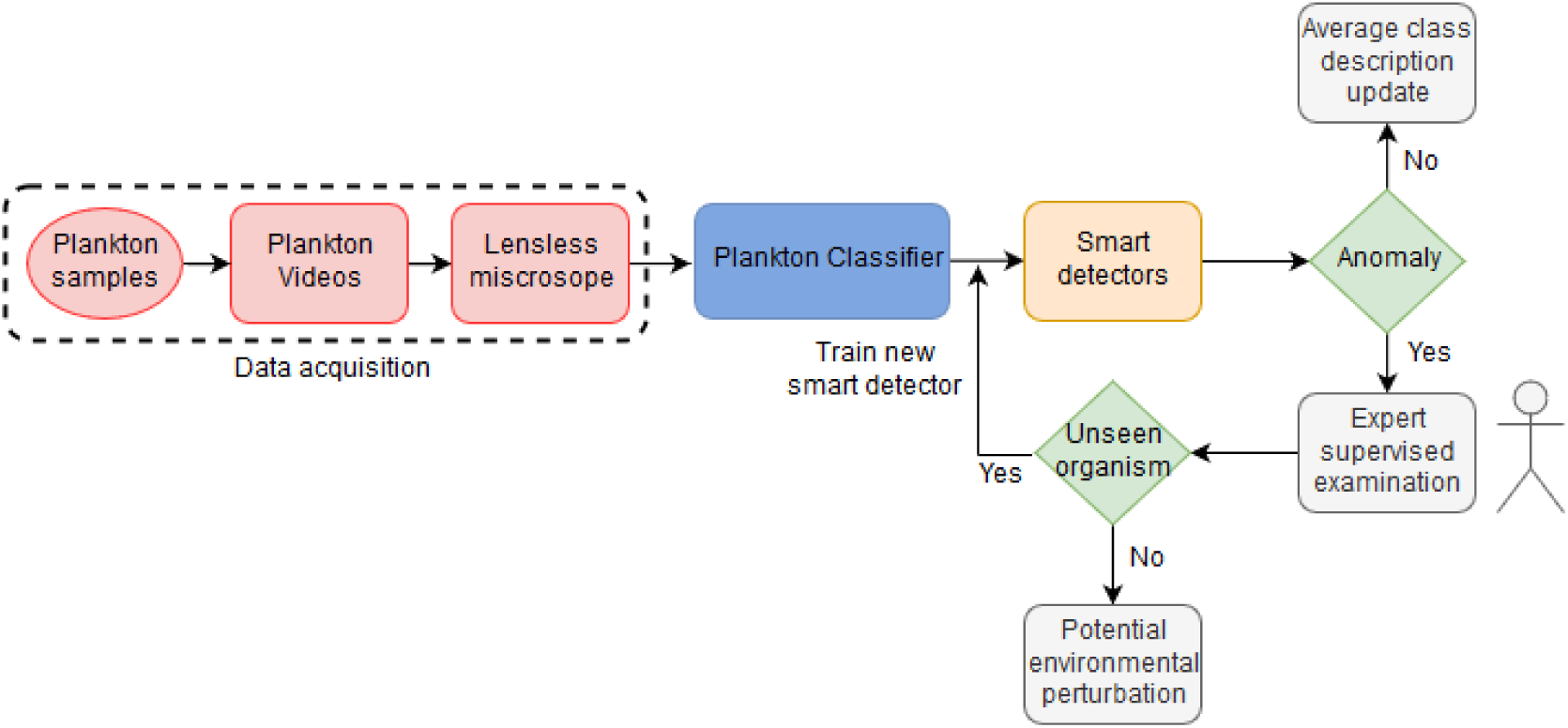
Proposed real-time smart environmental monitoring pipeline.

Finally, it is interesting to consider if such unsupervised approach can be utilized for different data types, thus widening the potential applicability and interest of the technique. While an extensive analysis of the performance of our pipeline on diverse set of data is beyond the scope of this work, it is worth commenting that the algorithms we use are general and pose no evident drawback to their application to other cell types. Particularly, the features our classifier uses to cluster the images do not include anything specific to plankton species (e.g. detection and estimation of number of flagella or other organelles.) Moreover, the proposed Deep Feature extraction method is even less dependent on the kind of data under study and may increase the applicability to other cell types. Thus, we expect the method to be potentially useful to other biological imaging fields.

## Material and methods

The proposed unsupervised pipeline (i.e., the plankton classifier) shown in Fig 1, consists of four modules: an **image processor, a feature extractor, an unsupervised partitioning module** and a **classification module**. In the following paragraphs we provide a description of the modules in more details, along with the methods considered and tested that led to our final design.

### Image Processing

Each video consists of ten seconds of color video (1920×1080) captured at 30 frames per second. Background subtraction is applied to each frame to detect the swimming plankton in the image. A contour detector is applied to the processed image to create a bounding box around each plankter. Because of instrument design, organisms can swim in and out of the field of view (FOV) during acquisition. Our algorithm automatically selects organisms which are fully contained inside the FOV by checking whether the bounding box touches the borders of the FOV. In this way, the images we obtain will be only of fully visible organisms. The resulting cropped image is then saved. From this collection of images, a training set of 640 images (500 training and 140 testing) is selected for each class. An image processor module for static images has also been implemented for benchmarking the plankton classifier on existing plankton datasets (e.g., the WHOI dataset; See Supporting Information for further details.).

### Feature Extraction

For each plankter image, 131 features are extracted from four categories: geometric (14), invariant moments (32), texture (67) and Fourier descriptors (10). Geometric features include area, eccentricity, rectangularity and other morphological descriptors, that have been used to distinguish plankton by shape and size [16]. The invariant Hu [28](7) and Zernike moments [29] (25) are widely used in shape representation, recognition and reconstruction. Texture based features encode the structural diversity of plankton. Fourier Descriptors (FD) are widely used in shape analysis as they encode both local fine-grained features (high frequency FD) and global shapes (low frequency FD). A full list of the features we have selected is reported in Table 1. These features span a 131-dimensional space, capturing the biological diversity of the acquired plankton images. Figs 3a and 3b demonstrate as an example, the discriminating power of the geometrical features for the ten evaluated species.

### Convolutional Neural Network (CNN) for deep features extraction

We implemented a deep CNN using eight convolutional layers and two fully connected layers, as described in Fig 6. We customized our architecture to be invariant with respect to rotation, similar to what has been done in [18]. Each input sample is rotated four times at multiples of 90 degrees, and all the tensors resulting from the features extraction module are concatenated and used to train the fully connected layers. The neural network has been trained for 60 epochs, using stochastic gradient descent with learning rate equal to 10^-5^, using data augmentation by means of translation, zooming, and rotation. It is worth noticing that the implemented rotational invariance module actually performs a data augmentation operation, and it is indeed useful when partial training data are available.

**Fig 6.**
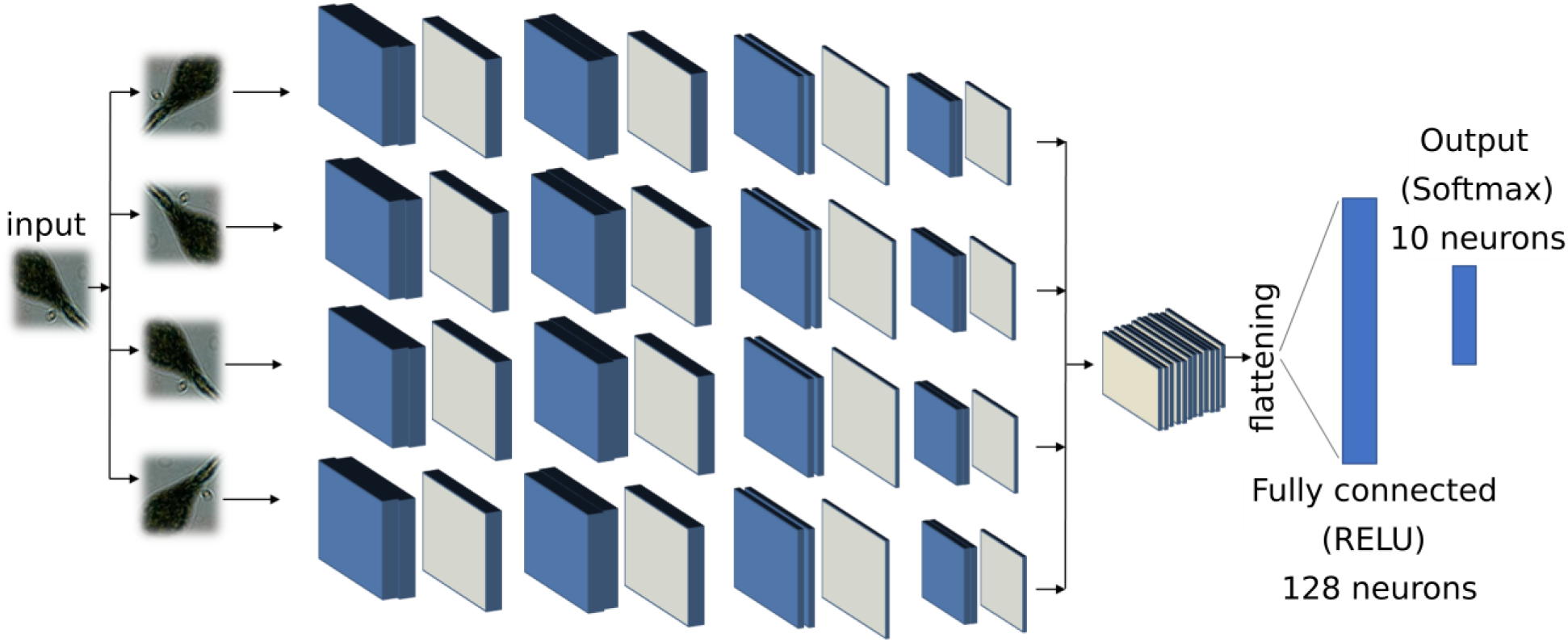
Deep features extraction. Deep CNN implemented for the purpose of deep features extraction. The blue layers represent convolutional layers, the grey ones represent a max pooling 2D operation. The fully connected layer with 128 neurons output has been used as feature set to the subsequent modules in our pipeline.

### Unsupervised Partitioning

#### Partition Entropy (PE)

The Partition Entropy (PE) coefficient is defined as:

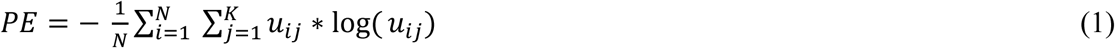

The coefficient is computed for every *j* in [0, K] and takes values in range [0, log(K)]. The estimated number of clusters is assigned to the index *j\** corresponding to the maximum PE value, PE(*j*\*). The lower the PE(*j\**), the higher the uncertainty of the clustering. We repeat this procedure ten times and obtain a distribution of *j\**. Finally, the estimation of the number of clusters *Z* is the mode of this distribution.

#### Clustering accuracy

Clustering accuracy is evaluated using purity:

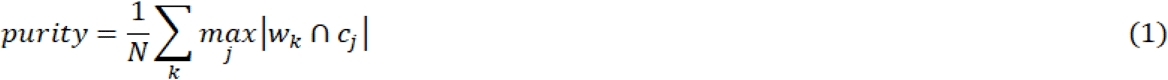

where the class *k* is associated to the cluster *j* with the highest number of occurrences. A purity value of one corresponds to clusters that perfectly overlap the ground truth. Purity decreases when samples belonging to the same class are split between different clusters, or when two or more clusters overlap with the same species. We have implemented a purity algorithm capable of checking for these occurrences and automatically adapt to the correct number of non-overlapping clusters (see Supporting Information).

### Classification algorithms

#### Random Forest

Random Forests (RF) is a popular ensemble learning method [30] used for classification and regression tasks, introduced in 2001 by Breiman. Random forests model providing estimators of either the Bayes classifier or the regression function. Basically, RF work building several binary decision trees using a bootstrap subset of samples coming from the learning sample and choosing randomly at each node a subset of features or explanatory variables [31]. Random forests are often used for classification of large set of observations. Each observation is given as input at each of the decision tree, which will output a predicted class. The model outputs the class that is the mode of the class output by individual trees [32].

Let us consider a set of observations *x*_1_, *x*_2_, …, *x*_n_, with *x* ∈ *R*^m^. The decision tree is designed as follows: we extract N times from the set of training observations (with replacement), for a each of the total number of decision tree. We specify the number of features *m*\* to consider for the tree growing, with *m*\* ≪ *m*. For each of the nodes in the tree, the algorithm randomly selects *m*\* features and calculates the best split for that node. The trees are only grown and not pruned (as in a normal tree classifier [33]. The split’s aim is to reduce the classification error at each branch. In detail, the algorithm considers an entropy-based measure trying to reduce the amount of entropy at each branch, selecting, with such a procedure, the best split. A possible choice is the Gini index:

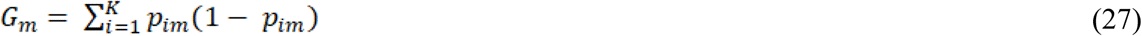

Where *G_m_* is the Gini Index for branch at level m in the decision tree, and *p_im_* is the proportion of observations assigned to class *i*. Minimizing *G_m_*, means to decrease the heterogeneity at each branch, i.e., a best split will correspond to a lower number of class in the children nodes. The algorithms continue in growing trees until convergence on the entropy-based on the generalization error [32].

#### Neural Networks

An artificial neural network (or multi-layer perceptron) is a massive parallel combination of single processing unit which can acquire knowledge from environment through a learning process and store the knowledge in its connections [23]. Classification is one of the most active research and application areas of neural networks. In this work we used an artificial neural network to build a classifier able to predict the species for each observation extracted using the shadow microscope. Fig. 2 shows the developed architecture. The network is very shallow, with two hidden layers of 40 neurons and an output layer with as much neurons as the number of species to classify. As reported in the main text of this manuscript, we used a training dataset with 10 species, thus the output layer is made up of k neurons, where k is the number of clusters obtained using the unsupervised clustering. As Fig 7 shows, the developed NN uses RELU activation function and dropout to reduce the overfitting. The network was trained using 200 epochs, Root mean square as an optimizer, a learning rate *λ* = 0,005 and categorical cross-entropy as loss function. The training requires 50 seconds on a MAC book PRO, core i7 – 2.9 GHz, solid state disk and 16 GB of RAM. The neural network has been implemented using KERAS, a powerful high-level neural network API running on top of TensorFlow.

**Fig 7.**
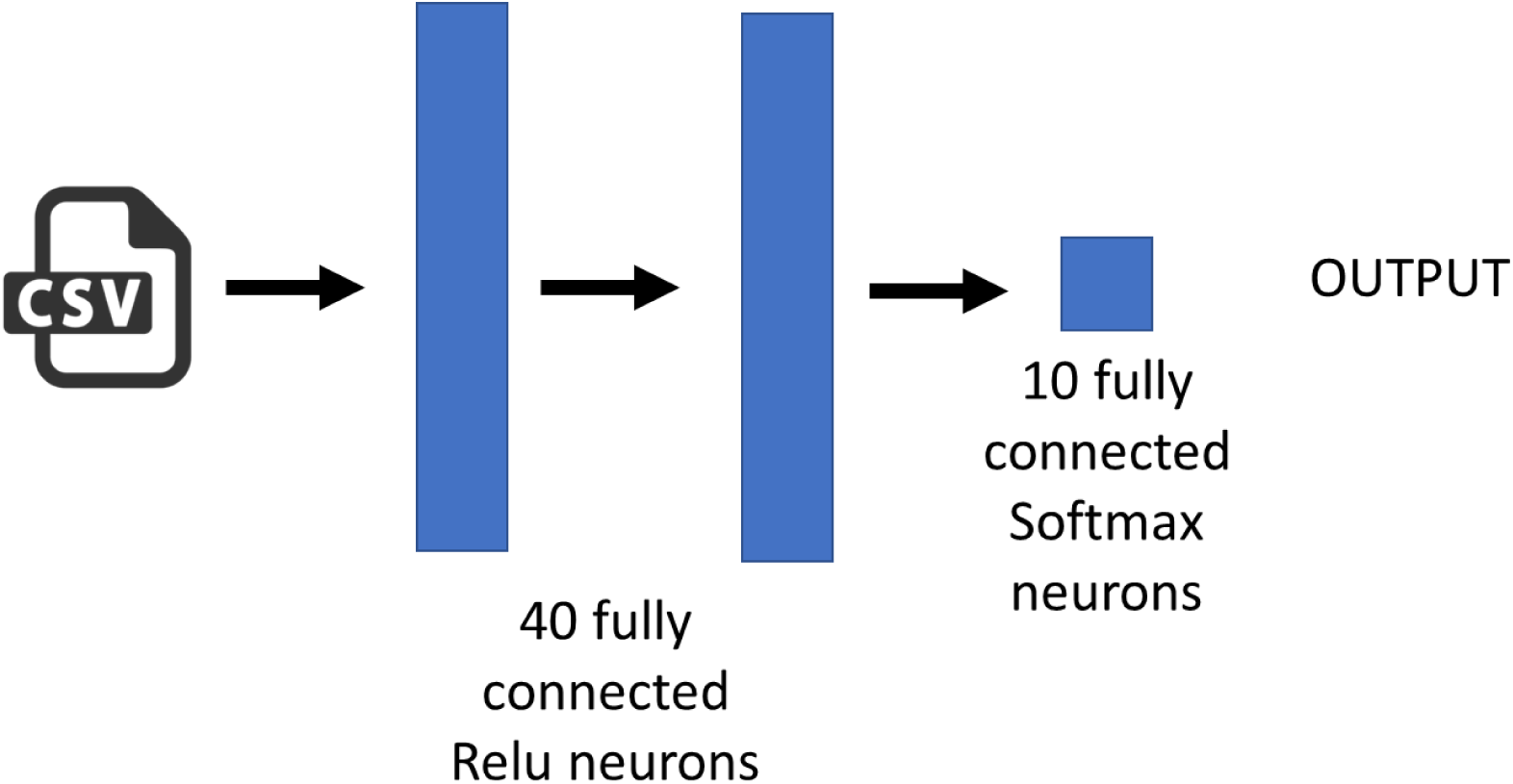
ANN architectures implemented for classification based on the extracted features.

### Anomaly Detection

#### One Class SVM

We adopted the one class SVM described by Scholpoff in [34]. Let us consider a set of N observations: 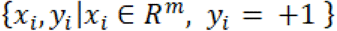. Where *x_i_* is a m-dimensional real vector and *y_i_* = +1 simply imply that the set contains normal observations belonging to a certain class. The one-class SVM is a classification algorithm returning a function which takes +1 in a “small” region capturing most of the data points, and −1 elsewhere. Let *ϕ* be a feature map that map our observations set *x_i_*, into an inner product space such as the inner product for the image of *ϕ* can be evaluated using some simple kernel:

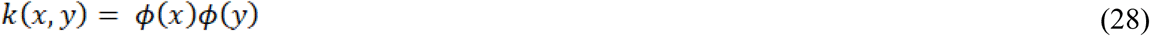

The strategy of the one class SVM is to map the data into the kernel space and separate the data from the origin with maximum margin, defining a hyperplane as:

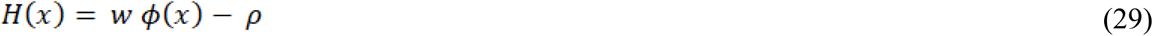

Meaning that we want to maximize the ratio 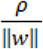, corresponding to the hyperplane’s distance from the origin. In order to solve this maximization problem, we have to solve a quadratic problem:

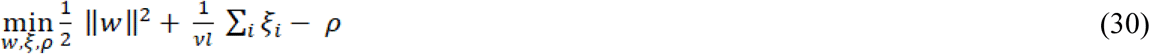

subject to 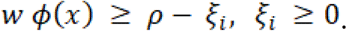

Where *ϕ*(*x*) is the feature mapping function that maps observations x into a feature space, *ξ_i_* is a slack variable for outlier that allows observations to fall on the other side of the hyperplane, 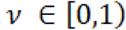 is a regularization parameter determining the bounding for the fractions of outliers and support vectors.

If *w* and *ρ* solve this problem, then the decision function:

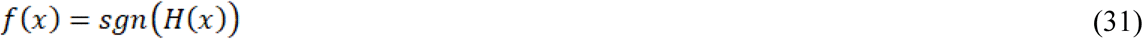

will be positive for most of the training observation, while w will be still small. The parameter influences the trade-off between the reported properties. To solve the quadratic form, we can use Lagrangian multipliers, obtaining:

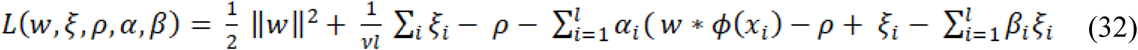

And set the derivatives with respect to w, *ξ* and *ρ* and expanding using the kernel expression yields:

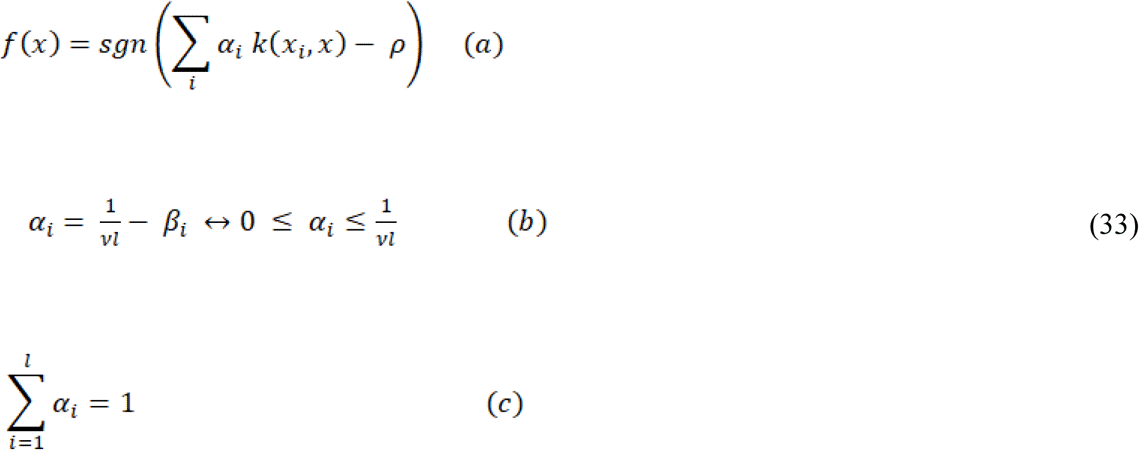

We used a Radial Basis Function kernel (RBF):

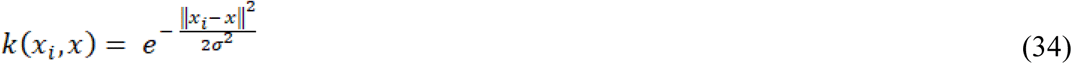

And then the original quadratic problem is solved substituting Eq. 16 into Eq. 15, yielding:

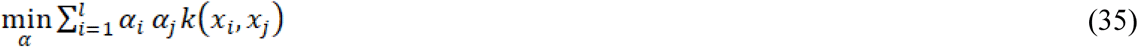

under the constraint of Eq. (16b) and (16c).

We finally use the support vectors to recover the parameter needed to compute the hyperplane:

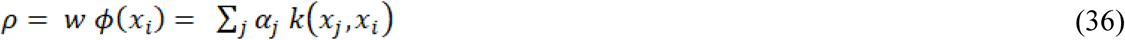

### DEC detectors

We designed a deep neural network that we named Delta-Enhanced Class (DEC) detector for the purpose of anomaly detection. The DEC detector’s architecture is represented in Fig 8, and shows a 2-neurons output, indicating that the sample is a member of the class or is an anomaly (i.e. not a member of the class). For each observation, we train such neural network with the actual features vector and extract randomly select a set of points from the training class in our dataset. For each of these selected points, we define a custom network layer (delta layer) that computes the difference in absolute value (as a vector, feature by feature) between the actual observation and the extracted random set. The vector of differences and the actual observations are used as inputs to the neural network (Fig 8), which assigns the proper weights to either one during training. The set of points to select is a hyperparameter which needs to be tuned. Through testing we determine that 25 points is the optimal tradeoff accuracy and computational cost.

**Fig 8.**
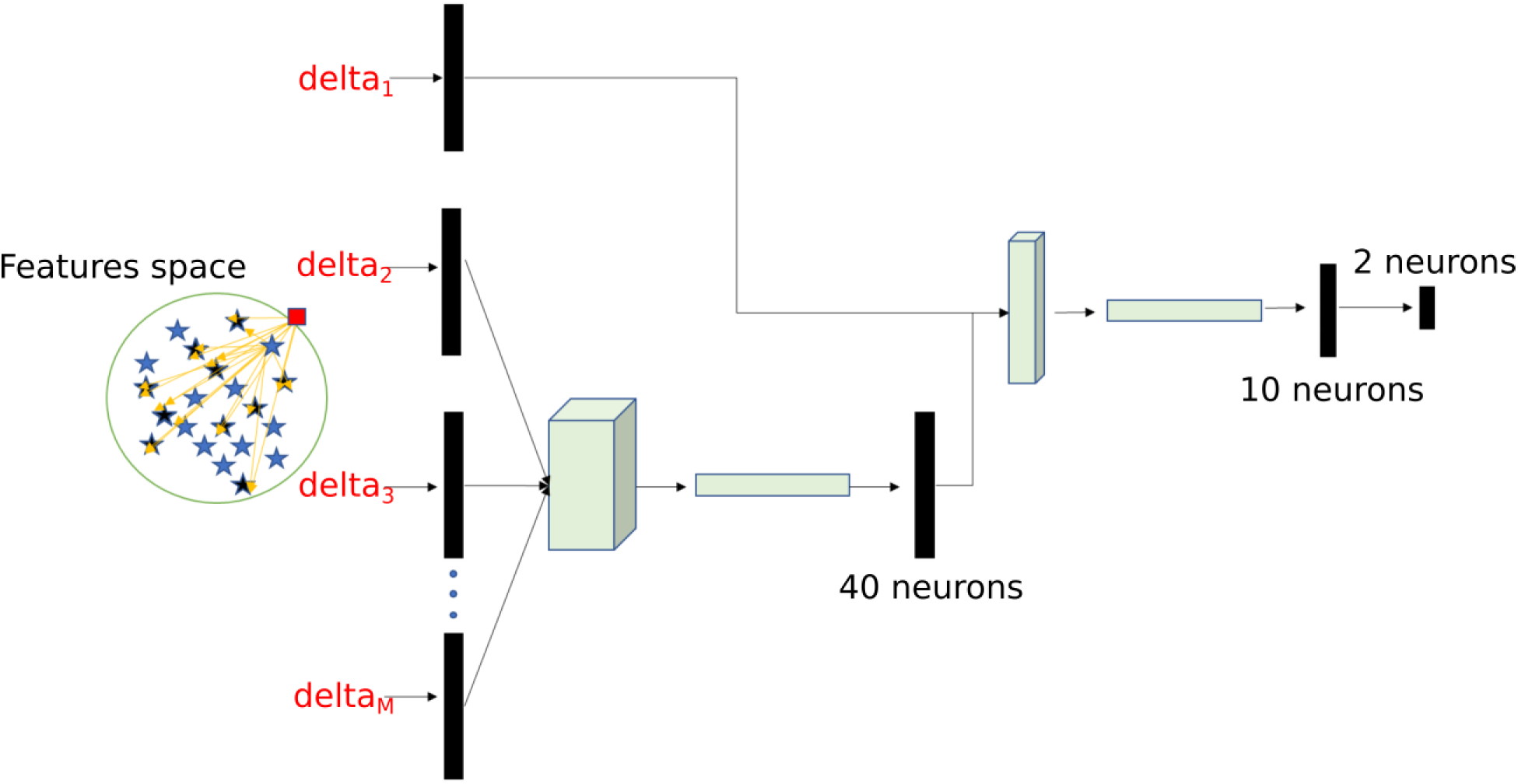
Schematic representation of DEC detector architecture.

### Code availability

The full source code accompanying this paper has been made available under EPL license at the following link: https://github.com/sbianco78/UnsupervisedPlanktonLearning.

## Supporting information

**S1 Data**. **The lensless microscope dataset and the dataset extracted from the WHOI used in this paper is available at the following link:** https://ibm.ent.box.com/s/8g2mp5knl2by7cv0ie0fx60mlb3rs6v3

**S1 Text. Supplementary Information include: S1**. Implemented detector to extract plankton images from the acquired videos **S2.** Evaluation of purity with respect to the number of samples using the lensless microscope dataset **S3.** Example images from the considered datasets **S4.** Example images from the considered datasets **S5.** *Estimated number of clusters adopting the partition coefficient* **S6.** *Local Binary Pattern computation.* **S7.** *Multi-dimensional representation for the Haralick subset of features* **S8.** *Multi-dimensional representation for the Hu-moments subset of features* **S9.** *Multi-dimensional representation for the features extracted from the gray values histogram* **S10.** *Multi-dimensional representation for the LBP subset of features* **S11.** *Multi-dimensional representation for the Fourier Descriptors subset of features* **S12.** *Multi-dimensional representation for the Zernike moments subset of features* **S13.** *Histogram reporting the normalized ranking score for the set of designed descriptors* **S14.** Schematic work flow describing how an observation is associated to the three possible outpus of the developed system: retraining class, anomaly or belonging to a trained class

**S1 Fig. Implemented detector to extract plankton images from the acquired videos.** The bounding box corresponding to the final detected contour is used to crop the plankton image.

**S2 Fig. Evaluation of purity with respect to the number of samples using the lensless microscope dataset.** The results are very accurate with number of images per sample higher or equal to 100. Using 50 images results in an overlap between two clusters (corresponding to the species Paramecium bursaria and Blepharisma americanuum), and in a decrease of the performances (light gray bar). The corrected purity algorithm introduced in this supplement (see Customized purity algorithm section), allows for a more accurate result (patterned bar).

**S3 Fig. Example images from the considered datasets. a-z13** WHOI dataset (names as they are labeled in the dataset) **z14-z23** lensless microscope dataset. **a** Ceratium **b** Chrysochromulina **c** Coscinodiscus **d** Dactyliosolen **e** Gyrodinium **f** Strombidium_morphotype1 **g** Dino30 **h** Euglena **i** Eucampia **j** Flagellate_sp3 **k** Pyramimonas_longicauda **l** Thalassionema **m** Delphineis **n** Pleurosigma **o** Chaetoceros_didymus_flagellate **p** Dictyocha **q** DactFragCerataul **r** Emiliania_huxleyi **s** Corethron **t** Kiteflagellates **u** Tintinnid **v** Dinobryon **w** Ephemera **x** Thalassiosira_dirty **y** Skeletonema **z** Pseudochattonella_farcimen **z0** Proterythropsis_sp **z1** Heterocapsa_triquetra **z2** Rhizosolenia **z3** Prorocentrum **z4** Pleurosigma **z5** Phaeocystis **z6** Laboea Strobila **z7** Katodinium_or_Torodinium **z8** Mesodinium_sp **z9** Paralia **z10** Guinardia_striata **z11** Asterionellopsis **z12** Amphidinium_sp **z13** Pennate_morphotype1 **z14** Blaepharisma Americanum **z15** Euplotes Eurystomus **z16** Spirostomum ambiguum **z17** Volvox **z18** Arcella Vulgaris **z19** Actinosphaerium Nucleofilum **z20** Dileptus **z21** Stentor Coeruleous **z22** Paramecium Bursaria **z23** Didinium nasutum.

**S4 Fig. Examples of species that are incorrectly assigned to the same cluster by our algorithm because of their morphological similarity in our feature space**. Similarity is intended from left to right **a** Proterythropsis_sp **b** Heterocapsa_triquetra **c** Amphidinium_sp **d** Pseudochattonella_farcimen **e** Gyrodinium **f** Prorocentrum

**S5 Fig. Estimated number of clusters adopting the partition coefficient. a** and the XIE-BENI index **b** as a function of sample size (species). The results are less precise if compared with the partition entropy (see fig 2e in the main text). However, both the algorithms can reconstruct correctly the number of clusters for subset of 3 species and 5 species. The number of clusters on the y axis is the distribution of ten runs on random subsets of all species. For example, for the leftmost box, 3 species have been randomly chosen from the lensless microscope database. This procedure is repeated ten times and the mode is then used as the estimated number of clusters.

**S6 Fig. Local Binary Pattern computation.**

**S7 Fig. Multi-dimensional representation for the Haralick subset of features. a** Andrew’s curve**. b** Parallel coordinates

**S8 Fig. Parallel coordinate for the Hu-moments subset of features. a** Andrew’s curve**. b** Parallel coordinates

**S9 Fig. Multi-dimensional representation for the features extracted from the gray values histogram. a** Andrew’s curve**. b** Parallel coordinates

**S10 Fig. Multi-dimensional representation for the LBP subset of features. a** Andrew’s curve**. b** Parallel coordinates

**S11 Fig. Multi-dimensional representation for the Fourier Descriptors subset of features. a** Andrew’s curve**. b** Parallel coordinates

**S12 Fig. Multi-dimensional representation for the Zernike moments subset of features. a** Andrew’s curve**. b** Parallel coordinates

**S13 Fig. Histogram reporting the normalized ranking score for the set of designed descriptors.**

**S14 Fig. Schematic work flow describing how an observation is associated to the three possible outputs of the developed system: retraining class, anomaly or belonging to a trained class**

**S1 Table. Computational time on raspberry pi for the analysis of one sample.** The standard deviation is computed among the objects contained into the 60 frames of the analyzed video.

## Acknowledgment

We thank Amanda K. Paulson and Aleksandar Godjoski for critical reading of the manuscript. We also thank all faculty and students in the National Science Foundation Center for Cellular Construction for discussion and critical feedback on the general idea and pipeline.

## Author contribution

**Conceptualization:** Vito Paolo Pastore, Simone Bianco.

**Data curation:** Vito Paolo Pastore, Simone Bianco.

**Funding acquisition:** Simone Bianco.

**Investigation:** Vito Paolo Pastore, Simone Bianco and Thomas Zimmerman.

**Methodology:** Vito Paolo Pastore, Sujoy K. Biswas, Thomas Zimmerman and Simone Bianco.

**Project administration:** Simone Bianco.

**Resources:** Simone Bianco.

**Software:** Vito Paolo Pastore

**Supervision:** Simone Bianco and Thomas Zimmerman.

**Validation:** Vito Paolo Pastore, Sujoy K. Biswas, Thomas Zimmerman and Simone Bianco.

**Visualization:** Vito Paolo Pastore

**Writing ± original draft:** Vito Paolo Pastore, Thomas Zimmerman and Simone Bianco.

**Writing ± review & editing:** Vito Paolo Pastore, Thomas Zimmerman and Simone Bianco

## REFERENCES

[1] M. J. Behrenfeld et al., “Biospheric primary production during an ENSO transition,” Science, vol. 291, no. 5513, pp. 2594–2597, Mar. 2001.

[2] A. Sournia, M.-J. Chrdtiennot-Dinet, and M. Ricard, “Marine phytoplankton: how many species in the world ocean?,” J. Plankton Res., vol. 13, no. 5, pp. 1093–1099, Jan. 1991.

[3] A. J. Richardson et al., “Using continuous plankton recorder data,” Prog. Oceanogr., vol. 68, no. 1, pp. 27–74, Jan. 2006.

[4] T. O. Fossum et al., “Toward adaptive robotic sampling of phytoplankton in the coastal ocean,” Sci. Robot., vol. 4, no. 27, p. eaav3041, Feb. 2019.

[5] T. G. Zimmerman and B. A. Smith, “Lensless Stereo Microscopic Imaging,” in ACM SIGGRAPH 2007 Emerging Technologies, New York, NY, USA, 2007.

[6] Heidi M. Sosik, Emily E. Peacock, Emily F. Brownlee, “Annotated Plankton Images - Data Set for Developing and Evaluating Classification Methods.”

[7] M. S. Schmid, C. Aubry, J. Grigor, and L. Fortier, “The LOKI underwater imaging system and an automatic identification model for the detection of zooplankton taxa in the Arctic Ocean,” Comput. Vis. Oceanogr., vol. 15–16, pp. 129–160, Apr. 2016.

[8] Culverhouse, P. F., Ellis, R. E., Simpson, R. G., Williams, R., Pierce, R. W., Turner, J. T., “Categorisation of five species of Cymatocylis (Tintinidae) by artificial neural network,” 1994, pp. 107:273–280.

[9] E. C. Orenstein and O. Beijbom, “Transfer Learning and Deep Feature Extraction for Planktonic Image Data Sets,” in 2017 IEEE Winter Conference on Applications of Computer Vision (WACV), 2017, pp. 1082–1088.

[10] Lumini, Alessandra & Nanni, Loris, “Deep learning and transfer learning features for plankton classification,” 2019.

[11] Qiao Hu and Cabell Davis, “Automatic plankton image recognition with co-occurrence matrices and Support Vector Machine,” Marine Ecology Progress Series, vol. 295, pp. 21–31, 2005.

[12] M. C. B. | D. of Oceanography, et al., “RAPID: Research on Automated Plankton Identification,” Oceanography, vol. 20, Jun. 2007.

[13] Vito P. Pastore, Thomas Zimmerman, Sujoy K. Biswas, and Simone Bianco, “Establishing the baseline for using plankton as biosensor,” presented at the Proc.SPIE, 2019, vol. 10881.

[14] Sujoy Kumar Biswas et al., “High throughput analysis of plankton morphology and dynamic,” presented at the Proc.SPIE, 2019, vol. 10881.

[15] J. Dai, R. Wang, H. Zheng, G. Ji, and X. Qiao, “ZooplanktoNet: Deep convolutional network for zooplankton classification,” 2016, pp. 1–6.

[16] H. M. Sosik and R. J. Olson, “Automated taxonomic classification of phytoplankton sampled with imaging-in-flow cytometry,” Limnol. Oceanogr. Methods, vol. 5, no. 6, pp. 204–216, 2007.

[17] M. B. Blaschko et al., “Automatic In Situ Identification of Plankton,” in 2005 Seventh IEEE Workshops on Applications of Computer Vision (WACV/MOTION’05) - Volume 1, 2005, vol. 1, pp. 79–86.

[18] S. Dieleman, J. De Fauw, and K. Kavukcuoglu, “Exploiting Cyclic Symmetry in Convolutional Neural Networks,” ArXiv E-Prints, p. arXiv:1602.02660, Feb. 2016.

[19] H. Zheng, R. Wang, Z. Yu, N. Wang, Z. Gu, and B. Zheng, “Automatic plankton image classification combining multiple view features via multiple kernel learning,” BMC Bioinformatics, vol. 18, no. 16, p. 570, Dec. 2017.

[20] A. Hughes, J. D. Mornin, S. K. Biswas, D. P. Bauer, S. Bianco, and Z. J. Gartner, “Quantius: Generic, high-fidelity human annotation of scientific images at 105-clicks-per-hour,” bioRxiv, p. 164087, Jul. 2017.

[21] D. A. Reynolds, “Gaussian Mixture Models,” in Encyclopedia of Biometrics, 2009.

[22] A. Romero, C. Gatta, and G. Camps-Valls, “Unsupervised Deep Feature Extraction for Remote Sensing Image Classification,” IEEE Trans. Geosci. Remote Sens., vol. 54, no. 3, pp. 1349–1362, Mar. 2016.

[23] S. Haykin, Neural Networks: A Comprehensive Foundation, 1st ed. Upper Saddle River, NJ, USA: Prentice Hall PTR, 1994.

[24] M. H. Bhuyan, D. K. Bhattacharyya, and J. K. Kalita, “Network Anomaly Detection: Methods, Systems and Tools,” IEEE Commun. Surv. Tutor., vol. 16, no. 1, pp. 303–336, First 2014.

[25] Thomas Zimmerman et al., “Stereo in-line holographic digital microscope,” presented at the Proc.SPIE, 2019, vol. 10883.

[26] B. Grindstaff, M. E. Mabry, P. D. Blischak, M. Quinn, and J. C. Pires, “Affordable Remote Monitoring of Plant Growth and Facilities using Raspberry Pi Computers,” bioRxiv, p. 586776, Jan. 2019.

[27] C. Scherer et al., The development of UK pelagic plankton indicators and targets for the MSFD. 2015.

[28] Z. Huang and J. Leng, “Analysis of Hu’s moment invariants on image scaling and rotation,” 2010 2nd Int. Conf. Comput. Eng. Technol., vol. 7, pp. V7-476–V7-480, 2010.

[29] Z. Yang and T. Fang, “On the Accuracy of Image Normalization by Zernike Moments,” Image Vis. Comput, vol. 28, no. 3, pp. 403–413, Mar. 2010.

[30] T. K. Ho, “Random decision forests,” in Document analysis and recognition, 1995., proceedings of the third international conference on, 1995, vol. 1, pp. 278–282.

[31] R. Genuer, J.-M. Poggi, and C. Tuleau, “Random Forests: some methodological insights,” ArXiv08113619 Stat, Nov. 2008.

[32] L. Breiman, “Random Forests,” Mach. Learn., vol. 45, no. 1, pp. 5–32, Oct. 2001.

[33] “Random forest algorithm for classification of multiwavelength data - IOPscience.” [Online]. Available: http://iopscience.iop.org/article/10.1088/1674-4527/9/2/011. [Accessed: 11-Nov-2018].

[34] B. Schölkopf, J. C. Platt, J. Shawe-Taylor, A. J. Smola, and R. C. Williamson, “Estimating the support of a high-dimensional distribution,” Neural Comput., vol. 13, no. 7, pp. 1443–1471, Jul. 2001.

