## Supplementary Information for "Annotation-free Learning of Plankton for Classification and Anomaly Detection"

**Supplementary Material**

#### Dataset

##### Lensless microscope dataset

We acquired 1-minute videos of 10 species of plankton (Carolina Biological Supply, Burlington, NC) using an in-house digital detector. The system used for acquisition employs the principles of a lensless microscope. Using a point light sources, shadows of the plankton are cast upon an image sensor. We use a CMOS 5-megapixel image sensor (OmniVision OV5647). Sample images for the acquired species are provided in Fig. 1. The microscope camera acquisition frequency is 30 frames/s. To extract plankton images, each frame was analyzed using a detector implemented in python with openCV. The detector is based on 5 different steps (see fig S1):

-median blurring;

-background subtraction to eliminate static objects (e.g., algae in our videos);

-binarization of the image;

-morphological closing;

-contour extraction.

The extracted contour, as well as the cropped plankton cell images, are used to extract a set of 131 morphological descriptors, representing the input data for the unsupervised learning pipeline.

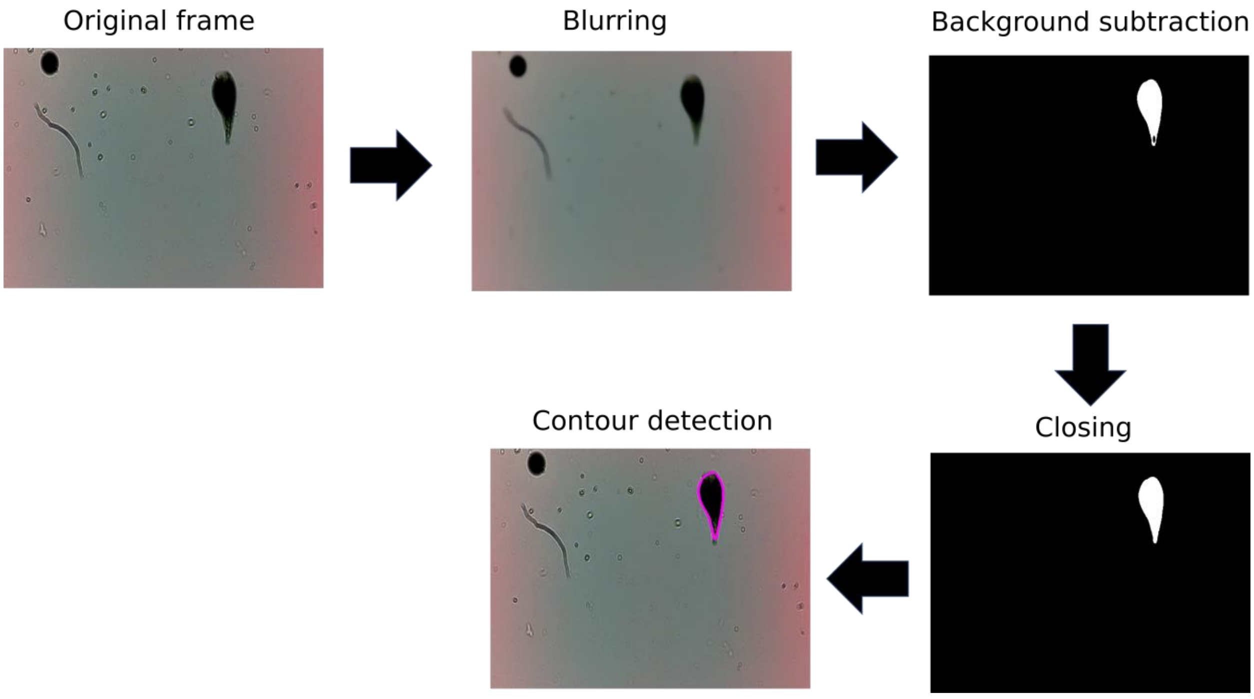


Fig **S1. Implemented detector to extract plankton images from the acquired videos.** The bounding box corresponding to the final detected contour is used to crop the plankton image.

#### Woods Hole Oceanographic Institution (WHOI) Dataset

The WHOI makes available a data set comprising of millions of still images of microscopic marine plankton, organized according to category labels provided by researchers at the Woods Hole Oceanographic Institution. These images were collected in situ by automated submersible imaging-in-flow cytometry with an instrument called Imaging FlowCytobot (IFCB), designed and built by WHOI. We have used this dataset as a benchmark to test the unsupervised partitioning approach we have developed. We started collecting the complete set of 103 categories acquired from 2011 to 2014. The WHOI dataset is strongly imbalanced, with number of images per classes varying from as low as ten images to several thousands. We have identified that 100 images is a low boundary for the size of our classes in order for the partitioning to be accurate (see fig S2). Thus, we eliminated all the classes which are represented in the database with less than 100 images, leaving us with 54 classes. The WHOI dataset has been acquired in the field, and then hand curated by experts. However, several of the annotated categories are macro categories containing more than one species of plankton (e.g., *pennate_on_diatoms* or *Chaetoceros_other*), for a total of 9 classes. We have decided to exclude these categories from the analysis. We have also excluded data which is not relevant to our work, like detritus or bubbles, for a total of 5 classes. After this trimming process, we obtained 40 species for which we were able to extract a random set of 100 images from the WHOI dataset. Fig S3 shows sample images extracted from the dataset for each of the selected species. We implemented a customized segmentation algorithm to make our pipeline compatible with static images, as the ones included into the WHOI dataset. The segmented image is the union between an adaptive gaussian thresholded image for detecting organism details, and a triangular thresholded image for reconstructing the organism body.

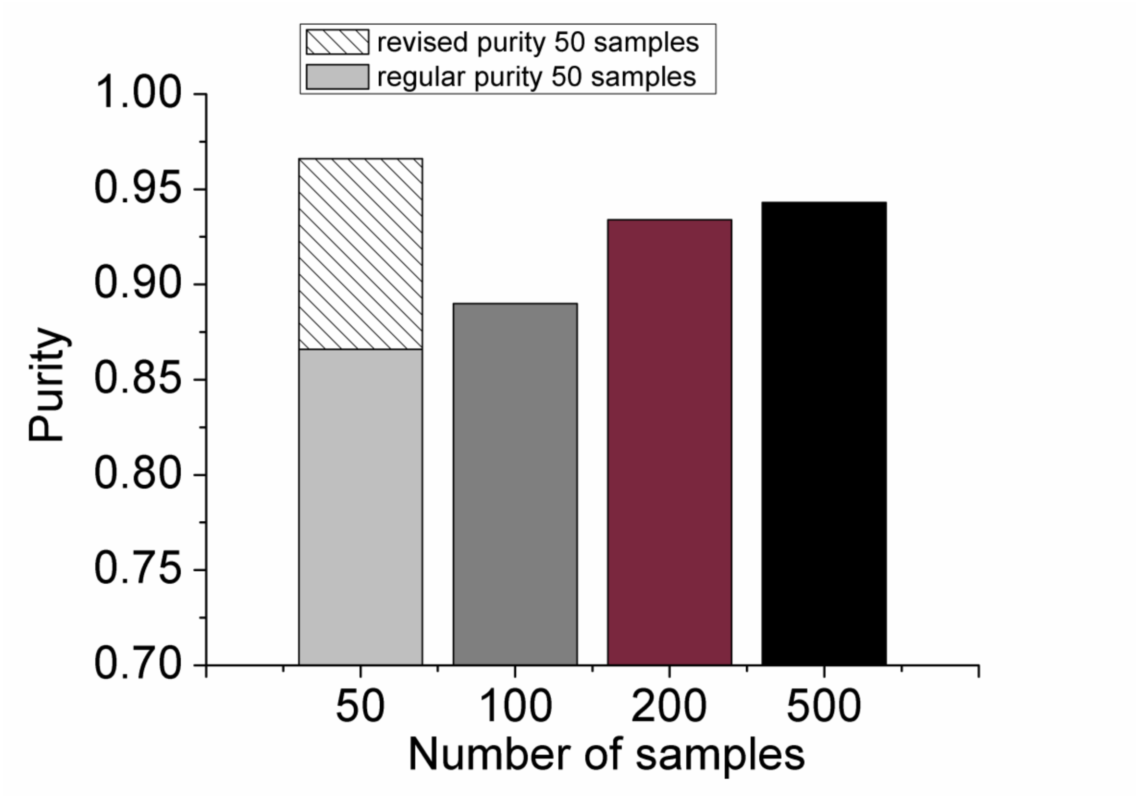


**Fig S2. Evaluation of purity with respect to the number of samples using the lensless microscope dataset.** The results are very accurate with number of images per sample higher or equal to 100. Using 50 images results in an overlap between two clusters (corresponding to the species Paramecium bursaria and Blepharisma americanuum), and in a decrease of the performances (light gray bar). The corrected purity algorithm introduced in this supplement (see Customized purity algorithm section), allows for a more accurate result (patterned bar).

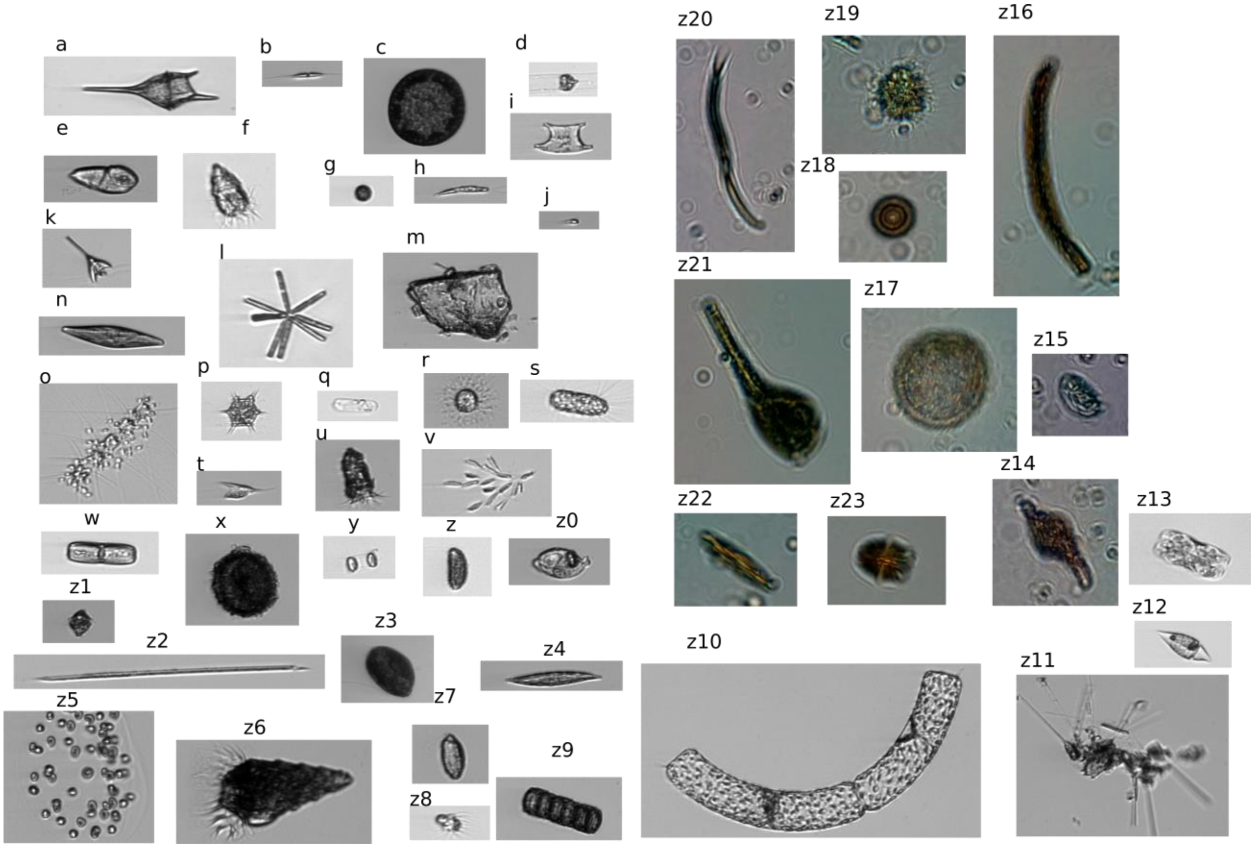


**Fig S3. Example images from the considered datasets. a-z13** WHOI dataset (names as they are labeled in the dataset) **z14-z23** lensless microscope dataset. **a** Ceratium **b** Chrysochromulina **c** Coscinodiscus **d** Dactyliosolen **e** Gyrodinium **f** Strombidium_morphotype1 **g** Dino30 **h** Euglena **i** Eucampia **j** Flagellate_sp3 **k** Pyramimonas_longicauda **l** Thalassionema **m** Delphineis **n** Pleurosigma **o** Chaetoceros_didymus_flagellate **p** Dictyocha **q** DactFragCerataul **r** Emiliania_huxleyi **s** Corethron **t** Kiteflagellates **u** Tintinnid **v** Dinobryon **w** Ephemera **x** Thalassiosira_dirty **y** Skeletonema **z** Pseudochattonella_farcimen **z0** Proterythropsis_sp **z1** Heterocapsa_triquetra **z2** Rhizosolenia **z3** Prorocentrum **z4** Pleurosigma **z5** Phaeocystis **z6** Laboea Strobila **z7** Katodinium_or_Torodinium **z8** Mesodinium_sp **z9** Paralia **z10** Guinardia_striata **z11** Asterionellopsis **z12** Amphidinium_sp **z13** Pennate_morphotype1 **z14** Blaepharisma Americanum **z15** Euplotes Eurystomus **z16** Spirostomum ambiguum **z17** Volvox **z18** Arcella Vulgaris **z19** Actinosphaerium Nucleofilum **z20** Dileptus **z21** Stentor Coeruleous **z22** Paramecium Bursaria **z23** Didinium nasutum.

### Clustering algorithms

#### Customized purity algorithm

We measure clustering accuracy using purity:

$$purity = \frac{1}{N}\sum_{k} \max_{j} |w_k \cap c_j| \qquad (1)$$

where the class k is associated to the cluster j with the highest number of occurrences. A purity value of one corresponds to clusters that overlaps perfectly to the ground truth. Purity decreases when samples belonging to the same class are split between different clusters, or when two or more clusters overlap with the same species. However, these two sources of error are very different. In fact, if two or more species are assigned to the same cluster, it means simply that they present very similar morphological features, and the clustering algorithm has joined them into the same cluster (see fig S4). If the morphological similarity is such that the overlapping species are fully assigned to the same cluster, the system is still capable of accurately representing the data, since the differentiation in the feature space of choice is, by construction, not be possible. We have customized the purity coefficient to take into account the possibility of these occurrences, by considering the overlap in the evaluation of the coefficient. Thus, if two species are morphologically indistinguishable and assigned to the same cluster, the index corresponding to total number of clusters is reduced by one in the calculation of the purity. On the other hand, when a species is assigned to multiple clusters, the error cannot be corrected since there would be no way to know that such error has occurred. We have observed errors of the first kind in the WHOI dataset. The number of potential misclassifications is correlated with the dimension of the training set, namely, the more images are available to the algorithm, the more accurate the separation in feature space, the lower the number of cluster overlaps. The maximum number of cluster misclassifcations is 5 for the subset of the WHOI we have considered, corresponding to using 50 images per class, in our analysis.

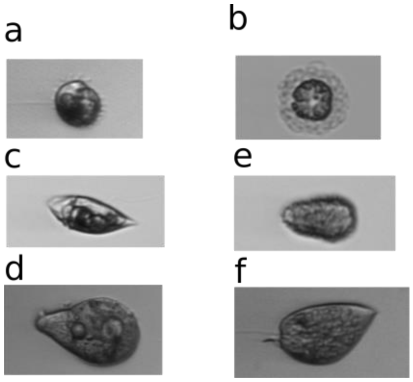


**Fig S4. Examples of species that are incorrectly assigned to the same cluster by our algorithm because of their morphological similarity in our feature space**. Similarity is intended from left to right **a** Proterythropsis_sp **b** Heterocapsa_triquetra **c** Amphidinium_sp **d** Pseudochattonella_farcimen **e** Gyrodinium **f** Prorocentrum

#### Estimation of the number of classes

We implemented three different methods to automatically determine the number of clusters, Z, to be used by the fuzzy k-means partitioning algorithm. The Partition Entropy (PE) is described in the main text, and its explanation will be excluded.

##### Partition Coefficient (PC)

The partition coefficient has been introduced in [1] and is defined as:

$$PC = \frac{1}{N}\sum_{i=1}^{N}\sum_{j=1}^{K} {u_{ij}}^2 \qquad (2)$$

The PC is evaluated for number of clusters greater or equal to 1, and the maximum value is adopted as an estimation of the total number of classes. The PC takes values in range [1/K,1]. The closer it is to 1/K, the smaller the separation between the analyzed data.

**Xie-Beni coefficient**

The Xie-Beni coefficient has been introduced in [2] and is defined as:

$$XB = \frac{\frac{1}{N}\sum_{i=1}^{C}\sigma_i^2}{{D_{min}}^2} \qquad (3)$$

Where:

$$\sigma_i^2 = \sum_{j=1}^{N} u_{ij}\left\|x_j - c_i\right\|^2$$

$x_j$ represents a N-dimensional feature vector that is used as input to the clustering algorithm. The Xie-Beni coefficient is also defined as compactness and separation coefficient. The logic of determination for the number of clusters relies in maximizing separation between clusters and compactness within the same cluster.

Both PC and Xie-Beni have been applied to the lensless microscope dataset, and have yielded incorrect results with respect to the PE (see fig S5).

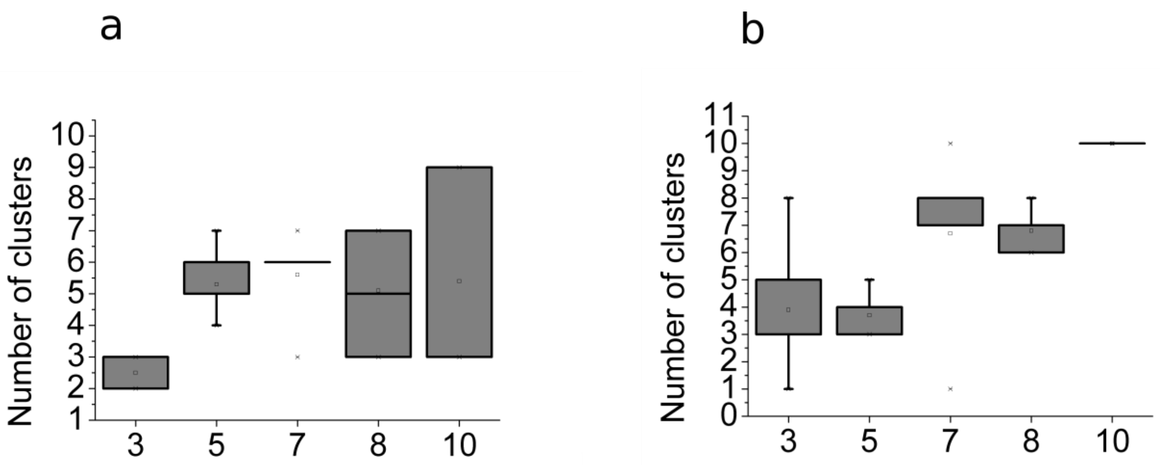


**Fig S5. Estimated number of clusters adopting the partition coefficient. a** and the XIE-BENI index **b** as a function of sample size (species). The results are less precise if compared with the partition entropy (see fig 2e in the main text). However, both the algorithms can reconstruct correctly the number of clusters for subset of 3 species and 5 species. The number of clusters on the y axis is the distribution of ten runs on random subsets of all species. For example, for the leftmost box, 3 species have been randomly chosen from the lensless microscope database. This procedure is repeated ten times and the mode is then used as the estimated number of clusters.

#### K-MEANS

K-mean is a is a clustering algorithm widely used in data mining, for clustering large sets of data. The term k-means was introduced for the first time by James MacQueen in [3]. The standard version of the algorithm, we are going to briefly describe in this section, is also referred as Lloyd’s algorithm and was introduced by Lloyd in [4]. The algorithm consists of two different steps. The first one consists in choosing the centroids ci for each of the k clusters. The second one consists in assigning each point to a cluster, minimizing an objective function. Let us consider a set of observations $x_1, x_2, \dots, x_n$, with $x \in R^m$. The k-mean algorithm will separate the observations into k partitions $P_1, P_2, \dots, P_n$, as to minimize the intra-partition variance:

$$\arg\min_P \sum_{i=1}^{k} \sum_{x \epsilon p_i} \|x - c_i\|^2 \quad (4)$$

After the first k centroids are choose, the algorithm assigns each observation x to the cluster correspondent to the nearest centroid (according to Euclidian distance). Then, the centroids are updated using the mean intra-cluster value, iterating the procedure until convergence (defined as no update in the centroids). The k-mean algorithm is a crispy clustering algorithm, i.e. there is a hard membership forcing a given point to belong exclusively to one cluster.

#### FUZZY K-MEANS

The fuzzy clustering is a set of algorithms that relaxes the crispy constraint of hard memberships, allowing each point to belong to two or more clusters. The fuzzy k-means was introduced in [5] and [6]. Given a set of observations $x_1, x_2, \dots, x_n$, with $x \in R^m$, the algorithm aims to minimize the following objective function:

$$\arg\min_P \sum_{i=1}^{N} \sum_{j=1}^{k} u^m{}_{ij} \left\|x_i - c_j\right\|^2, m \in [1, \infty) \quad (5)$$

The parameter *m* is defined fuzziness, $u^m{}_{ji}$ is the degree of membership for observation $x_i$ in cluster j, $c_j$ is the centroids for cluster j, ||*|| can be any norm to express similarity between observations and centroids. The centroids and the degree of memberships are updated according to Eq. (6) and (7):

$$u_{ij} = \frac{1}{\sum_{k=1}^{C}\left(\frac{\|x_i - c_j\|}{\|x_i - c_j\|}\right)^{\frac{2}{m-1}}} \tag{6}$$

$$c_j = \frac{\sum_{i=1}^{N} {u^m}_{ij} x_i}{\sum_{i=1}^{N} {u^m}_{ij}} \tag{7}$$

The algorithm converges to a saddle point for the objective function in Eq. (2), when there is a maximum change in the degree of membership lower than a given tolerance $\in$.

#### MIXTURES OF GAUSSIAN MODELS

Gaussian Mixture Model (GMM) is a parametric statistical model assuming data are generated from a linear combination of normal gaussian sources[7]. Given a set of observations $x_1, x_2, \dots, x_n$, with $x \in R^m$, the GMM model associates each observation xi with a probability p(xi) according to equation (8):

$$p(x_i|\Theta) = \sum_{i=1}^{K} \vartheta_i \, \mathcal{N}(x_i \,|\mu_i, \Sigma_i) \tag{8}$$

Where $\mathcal{N}(*)$ is a normal multivariate distribution with mean $\mu_i$ and covariance $\Sigma_i$, K is the number of gaussians distribution to consider, which corresponds to the number of clusters to partition the observations in. $\vartheta_i$ are the coefficient of the gaussians linear combination, with the constraint $\vartheta_i \geq 0$ and $\sum_{i=1}^{K} \vartheta_i = 1$. The complete set of parameters $\Theta$ is equal to the vector $(\vartheta_1, \vartheta_2, \ldots, \vartheta_K, \mu_1, \mu_{i2}, \ldots, \mu_K, , \Sigma_1, , \Sigma_2, \ldots, , \Sigma_K)$. To estimate $\Theta$, an Expectation-Maximization (EM) approach is adopted. Starting from an initial value for $\Theta$ (e.g., random), the E-step basically consists in computing the *N x K* matrix of the degree of membership for each observation $x_{\mathrm{i}}$ in cluster *j* as:

$$w_{ij} = \frac{\vartheta_j \, \mathcal{N}(x_i \,|\mu_j, \Sigma_j)}{\sum_{k=1}^{K} \vartheta_k \, \mathcal{N}(x_k \,|\mu_k, \Sigma_k)} \tag{9}$$

The degree of memberships satisfies the condition: $\sum_{j=1}^{K} w_{ij} = 1$. The M-step consists in update the parameter set $\Theta$ according to the estimated degree of membership matrix, according to Eq. 10, 11 and 12:

$$\vartheta_j = \frac{\sum_{i=1}^{N} w_{ij}}{N} \tag{10}$$

$$\mu_j = \frac{1}{\sum_{i=1}^{N} w_{ij}} * \sum_{i=1}^{N} w_{ij} \, x_i \tag{11}$$

$$\Sigma_j = \frac{1}{\sum_{i=1}^{N} w_{ij}} * \sum_{i=1}^{N} w_{ij} \, (x_i - \mu_j) * (x_j - \mu_j)^T \tag{12}$$

the two-step process is iterated until convergence, defined as a change in the log-likelihood for the space of parameter lower than a determined tolerance $\in$.

#### Shape descriptors

##### Hu moments

The Hu moments descriptors are introduced in[8], where six absolute orthogonal invariants and one skew orthogonal invariant from the normalized image central moments are derived. The two-dimensional image moments of order (*p* + *q*) are defined as follows:

$$m_{pq} = \int_{-\infty}^{+\infty} \int_{-\infty}^{+\infty} x^p y^q \, f(x,y) dp dq, \qquad p,q = 0,1,2 \ldots \qquad (13)$$

The moment $m_{pq}$ is uniquely determined if f (x, y) is a continuous piecewise bounded function. The moments defined in Eq. (10) are not invariant with respect to rotation, translation and scaling operations. To reach invariance with respect to rotation, it is possible to consider the central moments:

$$\mu_{pq} = \int_{-\infty}^{+\infty} \int_{-\infty}^{+\infty} (x - \bar{x})^p (y - \bar{y})^q \, f(x,y) dp dq, \qquad p,q = 0,1,2 \ldots \qquad (14)$$

Where $\bar{x} = \frac{m_{10}}{m_{00}}$ and $\bar{y} = \frac{m_{01}}{m_{00}}$ and $(\bar{x}, \bar{y})$ is the centroid of the image, f (x, y).

Starting from the translation invariant central moments, scaling invariance is obtained by introducing the normalized central moments for the image f (x, y):

$$\eta_{pq} = \frac{\mu_{pq}}{{\mu_{00}}^{\alpha}}, \quad \alpha = \frac{p+q}{2} + 1 \tag{15}$$

Starting from the normalized central moments, Hu derived 7 invariant moments:

$$\phi_1 = \eta_{20} + \eta_{02}$$

$$\phi_2 = (\eta_{20} - \eta_{02})^2 + 4\,\eta_{11}$$

$$\phi_3 = (\eta_{30} - 3\eta_{12})^2 + (\eta_{03} - 3\eta_{21})^2$$

$$\phi_4 = (\eta_{30} + \eta_{12})^2 + (\eta_{03} + \eta_{21})^2$$

$$\phi_5 = (3\eta_{30} - 3\eta_{12})(\eta_{30} + \eta_{12})[(\eta_{30} + \eta_{12})^2 - 3(\eta_{21} + \eta_{03})^2] + (3\eta_{21} - \eta_{03})(\eta_{21} + \eta_{03})[3(\eta_{30} + \eta_{12})^2 - (\eta_{21} + \eta_{03})^2$$

$$\phi_6 = (\eta_{20} - \eta_{12})[(\eta_{30} + \eta_{12})^2 - (\eta_{21} + \eta_{03})^2] + 4\eta_{11}(\eta_{30} + \eta_{12})(\eta_{21} + \eta_{03})$$

$$\phi_7 = (3\eta_{21} - 3\eta_{03})(\eta_{30} + \eta_{12})[(\eta_{30} + \eta_{12})^2 - 3(\eta_{21} + \eta_{03})^2] + (3\eta_{12} - \eta_{30})(\eta_{21} + \eta_{03})[3(\eta_{30} + \eta_{12})^2 - (\eta_{21} + \eta_{30})^2$$

(16)

Hu moments are invariant with respect to rotation, scaling and translation.

#### Zernike moments

The Zernike moments[9] are a set of complexes, orthogonal polynomials defined over the interior of the unit circle $x^2 + y^2 = 1$. The two-dimensional Zernike moment for order (p, q) is defined as:

$$A_{mn} = \frac{m+1}{\pi} \iint f(x,y)\ V_{mn}(x,y)^* \, dx\, dy \tag{17}$$

With $m = 0, 1, 2, \ldots, \infty$, * denotes the complex conjugate, f(x, y) is the function to describe and $n$ is an integer where m-$|n|$ is even and $|n|$ is lower or equal to m and $A_{mn}{}^* = A_{m,-n}$.

$V_{mn}(x,y)$ orthogonal polynomial can be expressed in polar coordinate as:

$$V_{mn}(\rho,\theta) = R_{mn}(\rho)\ e^{jn\theta}$$

$$R_{mn}(\rho) = \sum_{s=0}^{\frac{m-|n|}{2}} (-1)^s F(m,n,s,\rho)$$

$$F(m,n,s,\rho) = \frac{(m-s)!}{s!\left(\frac{m+|n|}{2}-s\right)!\left(\frac{m-|n|}{2}-s\right)!}\rho^{m-2s}$$

(18)

Zernike moments show robustness to noise, expression efficiency and fast computation. We implemented a customized version of Zernike moments starting from [10].

#### Gray Scale Co-occurrence Matrix (GSCM) and Haralick features

The GSCM is one of the earliest methods for texture feature extraction proposed by Haralick et.al. back in 1973[11]. It describes the distribution of gray scale values in image by counting the occurrence of each of the possible sequence of intensity values between two pixels, with an offset ($d_x$, $d_y$). Given an image I with size (N x N), the GSCM is defined as:

$$p(i,j) = \sum\sum\begin{cases}1, if\ (I(x,y) = i)\ and\ I(x + d_x, y + d_y) = j \\ 0, otherwise\end{cases} \quad (19)$$

In its original definition, the GSCM is not invariant with respect to rotations. To reach a degree of rotational invariance it is possible to consider the offset ($d_x$, $d_y$) at multiple of 45 degrees, computing the resulting average GSCM.

Haralick introduced 14 features to be extracted from the GSCM. Considering p (i, j) the entrance *(i, j)* of the gray scale co-occurrence matrix, $N_g$ the number of the image's gray levels, and $p_x(i)$ and $p_y(j)$ the marginal probabilities:

$$p_x\,(i) = \sum_{j=1}^{Ng} p(i,j)$$

$$p_y\,(i) = \sum_{i=1}^{Ng} p(i,j) \qquad (20)$$

And $\mu$ and σ are the average value and the standard deviation of the marginal probabilities, respectively. In the following section, we will define the 13 Haralick features implemented and used for our application:

$$f_1 = \sum_{i=1}^{N_g} \sum_{j=1}^{N_g} p\,(i,j)^2$$

$$f_2 = \sum_{n=0}^{N_g-1} n^2 \sum_{i=1}^{N_g} \sum_{j=1}^{N_g} p(i,j)$$

$$f_3 = \sum_{i=1}^{N_g} \sum_{j=1}^{N_g} \frac{(i - \mu_y)(\,i - \mu_y)p(i,j)}{\sigma_x \sigma_y}$$

$$f_4 = \sum_{i=1}^{N_g} \sum_{j=1}^{N_g} (i - \mu)^2 p(i,j)$$

$$f_5 = \sum_{i=1}^{N_g} \sum_{j=1}^{N_g} \frac{p(i,j)}{1 + (i-j)^2}$$

$$f_6 = \sum_{k=2}^{2N_g} k \, p_{x+y}(k)$$

$$f_7 = \sum_{k=2}^{2N_g} (k - f_6)^2 \, p_{x+y}(k)$$

$$f_8 = -\sum_{k=2}^{2N_g} p_{x+y}(k) \log p_{x+y}(k)$$

$$f_9 = -\sum_{i=1}^{N_g} \sum_{j=1}^{N_g} p(i,j) \, \log p(i,j)$$

$$f_{10} = -\sum_{k=0}^{N_g-1} [k - \sum_{l=0}^{N_g-1} l \, p_{x-y}(l)]^2 \, p_{x-y}(k)$$

$$f_{11} = \sum_{k=0}^{N_g-1} p_{x-y}(k) \log p_{x-y}(k)$$

$$f_{12} = \frac{f_9 - Hxy1}{\max(Hx, Hy)}$$

$$f_{13} = \left(1 - e^{\sqrt{[-2(Hxy2 - f_9]}}\right) \quad (21)$$

where Hxy1 and Hxy2 are defined as:

$$Hxy1 = -\sum_{i=1}^{N_g}\sum_{j=1}^{N_g} p(i,j) \log[p_x(i)p_y(j)]$$

$$xy1 = -\sum_{i=1}^{N_g}\sum_{j=1}^{N_g} p(i,j) \log[p_x(i)p_y(j)]$$

$$Hxy1 = -\sum_{i=1}^{N_g}\sum_{j=1}^{N_g} p_x(i)p_y(j) \log[p_x(i)p_y(j)] \quad (22)$$

#### Local Binary Pattern (LBP)

LBP[12] are grey-scale and rotation invariant texture operators, widely used in the last years in several application of computer vision and image processing. LBP summarizes structures of the image comparing each pixel to its neighborhood[13]. Given a pixel I(i, j) in an image I, we consider a window of size k x k centered in (i, j). The intensity value of the pixel (i, j) is subtracted from all the pixels inside window, binarizing the resulting differences: each positive value is set to one, each negative (or null) value is set to zero. The resulting matrix is used to build a binary code concatenating each digit starting from the one on the up left, in clockwise sense. The built code is referred as local binary pattern (see fig S6). It is possible to use a subset of $2^p$ LBP to describe the texture of an image.

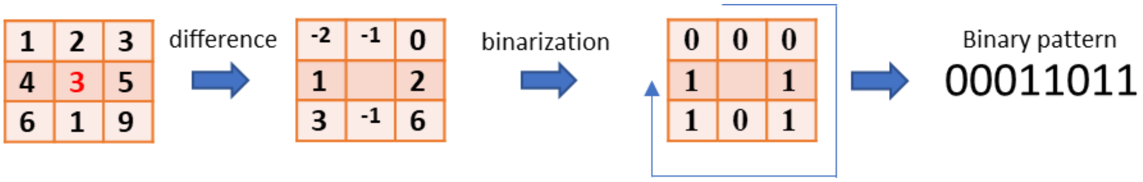


**Fig S6. Local Binary Pattern computation.**

#### Shape signatures and Fourier Descriptors (FD)

A shape signature[14] is a contour-based local representation of shape features, characterized by sensitivity to noise and not robustness. It has been showed that using shape signatures for computing FD improve the accuracy of the representation. A shape signature is a 1-D function representing 2D contour of an image. In our work, we used the centroid distance:

$$z(t) = \sqrt{(x(t) - x_c)^2 * (y(t) - y_c)^2} \quad (23)$$

Where $(x_c, y_c)$ is the image's centroid. The centroid distance is invariant with respect to translation.

Fourier transformation of the shape signature has been widely used for shape analysis and retrieval[15].

We used the set of points defining the contour of the analyzed shape provided by OpenCV to compute the FD. In detail, after computing the centroid distance signature, we evaluate the Fast Fourier Transform (FFT). Given our shape signature z(t), the FFT of z(t) is:

$$u_n = \frac{1}{N}\sum_{t=0}^{N-1} z(t)e^{\frac{-j2\pi nt}{N}}, n = 0,1, \dots, N-1 \qquad (24)$$

To get the rotation invariance, only the real part of the FFT[z(t)] is considered. A normalization with respect to the first Fourier coefficient is used to get scaling invariance:

$$FD = \left(\frac{FD_1}{FD_0}, \frac{FD_2}{FD_0}, \dots, \frac{FD_{N-1}}{FD_0}\right) \qquad (25)$$

It has been proved that the first ten FD can be enough to accurately describe image's shapes for retrieval and classification[14].

#### Multidimensional plot

A major problem for graphical representation of multivariate data is the dimensionality. In this manuscript, we exploited two different methods for visualizing our multidimensional set of features: parallel coordinates and Andrew’s curve.

#### PARALLEL COORDINATES

Parallel coordinates are a method of graphical representation of multidimensional data. It was introduced in [16], and consists in associating a N-dimensional data with N vertical parallel axis. Each observation will be represented by a multiline whose vertex on the j-th vertical axes represents the *j-th* coordinate for the observation, that is the specific value of the observation, among dimension *j*.

#### ANDREW’S CURVE

Andrew’s curve [17] provides visualization of multidimensional data by mapping them into a function, defined as:

$$f_x(\theta) = \frac{x_1}{\sqrt{2}} + x_2 \sin\theta + x_3 \cos\theta + x_4 \sin 2\theta + x_5 \sin 2\theta + \cdots \qquad (26)$$

It has been proved that Andrew's curves preserve mean, distance and variance of the multidimensional data, meaning that close Andrew's curves correspond to close data observations.

#### PLANKTON SPECIES MORPHOLOGICAL FEATURES SPACE

As reported in the main text of this manuscript, after a preprocessing detection step, we perform feature selection and extraction, which results in 131 features. Features were selected according to four characteristic classes, each shown to perform an efficient unsupervised partitioning and subsequent classification. Here we report a list of the designed features: geometric features (14 features), feature based on invariant moments (32 features), including Hu (7) and Zernike (25, up to order 5) moments, texture based features (67 features), including Haralick features, extracted from the grey scale co-occurrence matrix and the Local Binary Patterns (LBP), and, finally, the Fourier Descriptors (10 features). A full table of features is reported in the main text.

This set of features spans a 131-dimensional space, whose shape represents the projection of the biological diversity of the collection of organisms. In this section we report a graphical representation for each subset of features, previously described. For every subset of features, it is possible to appreciate the separation between the 10-species induced by the designed descriptor (see figs S7-S14)

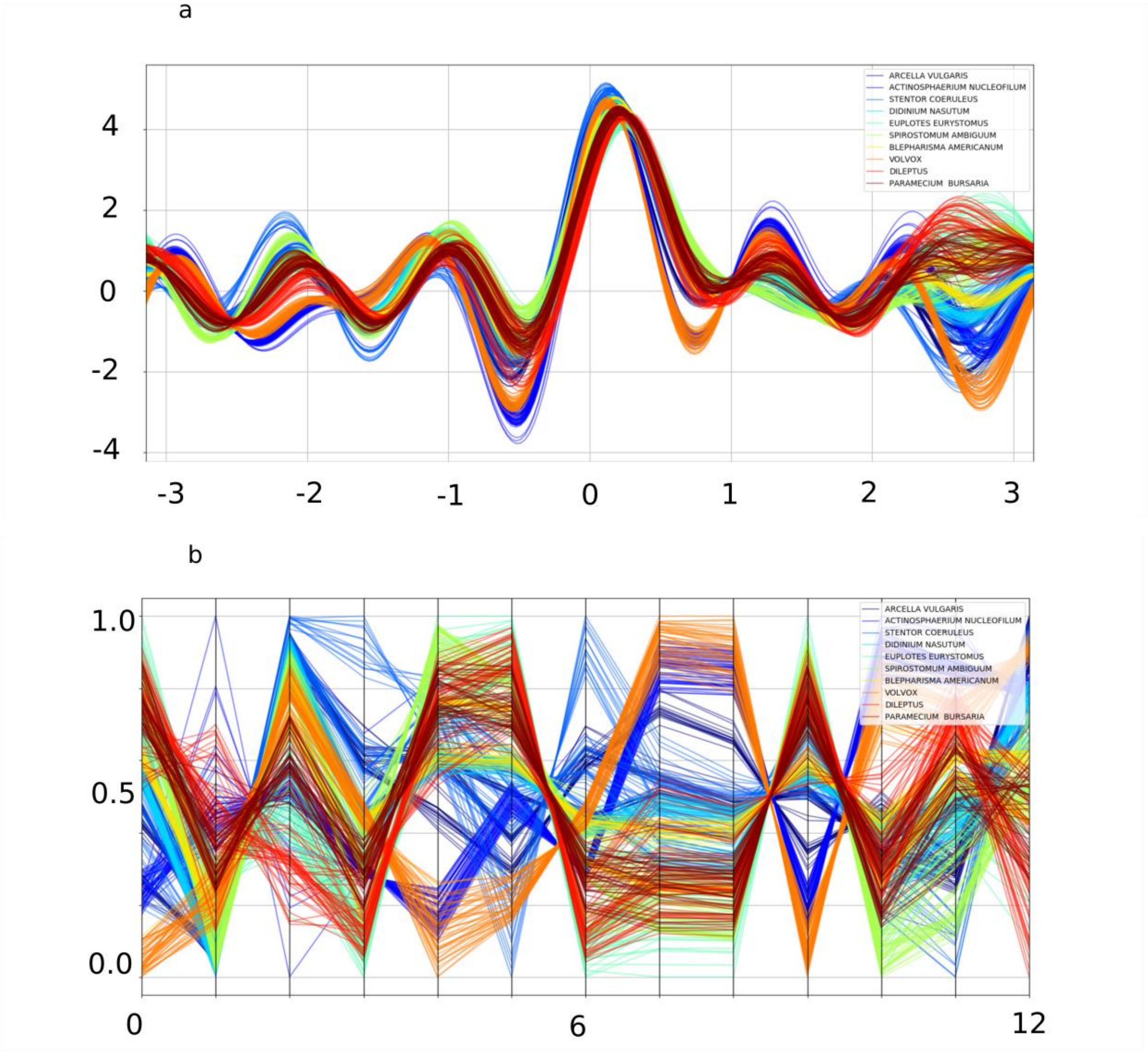


**Fig S7. Multi-dimensional representation for the Haralick subset of features. a** Andrew's curve**. b** Parallel coordinates

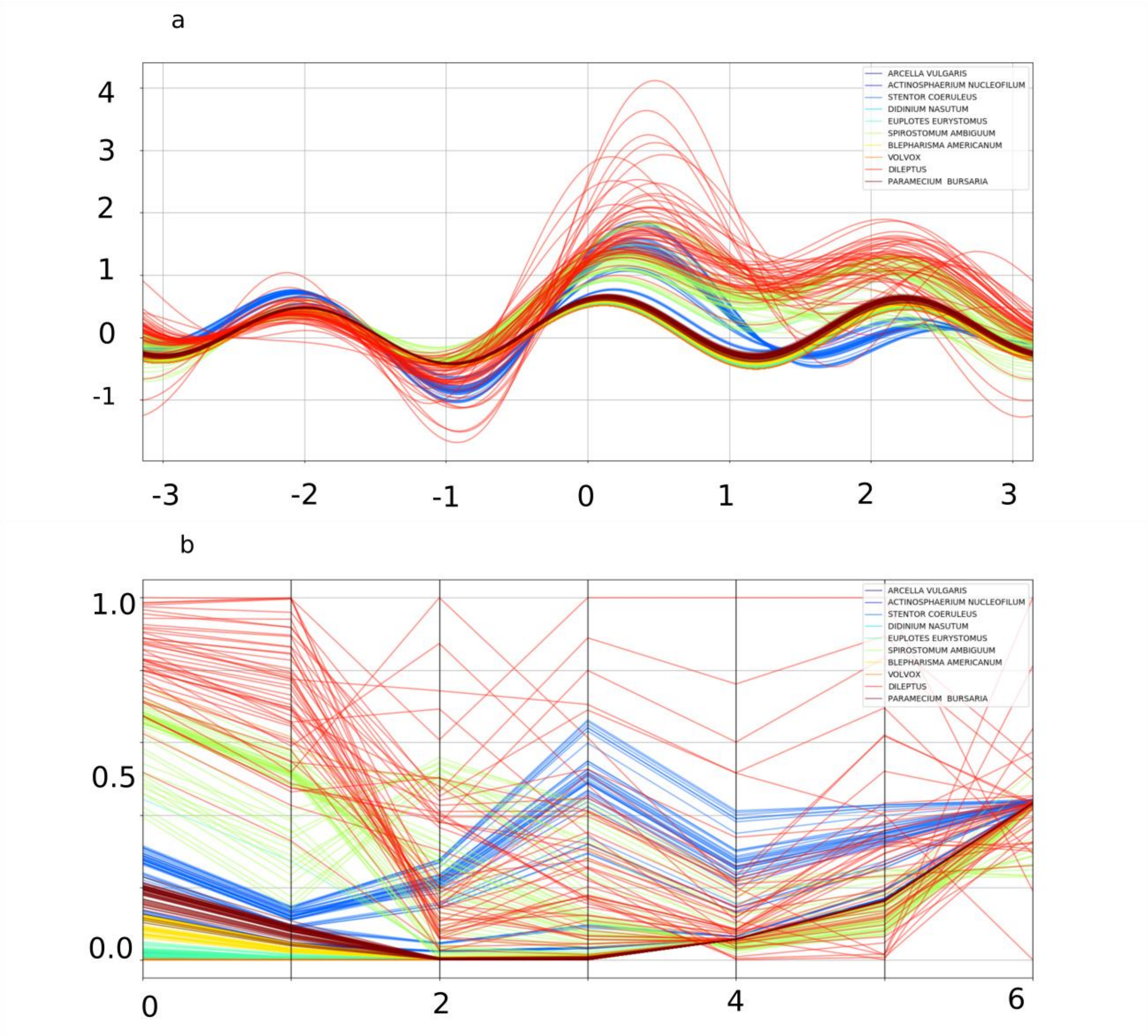


**Fig S8. Multi-dimensional representation for the Hu-moments subset of features. a** Andrew's curve**. b** Parallel coordinates

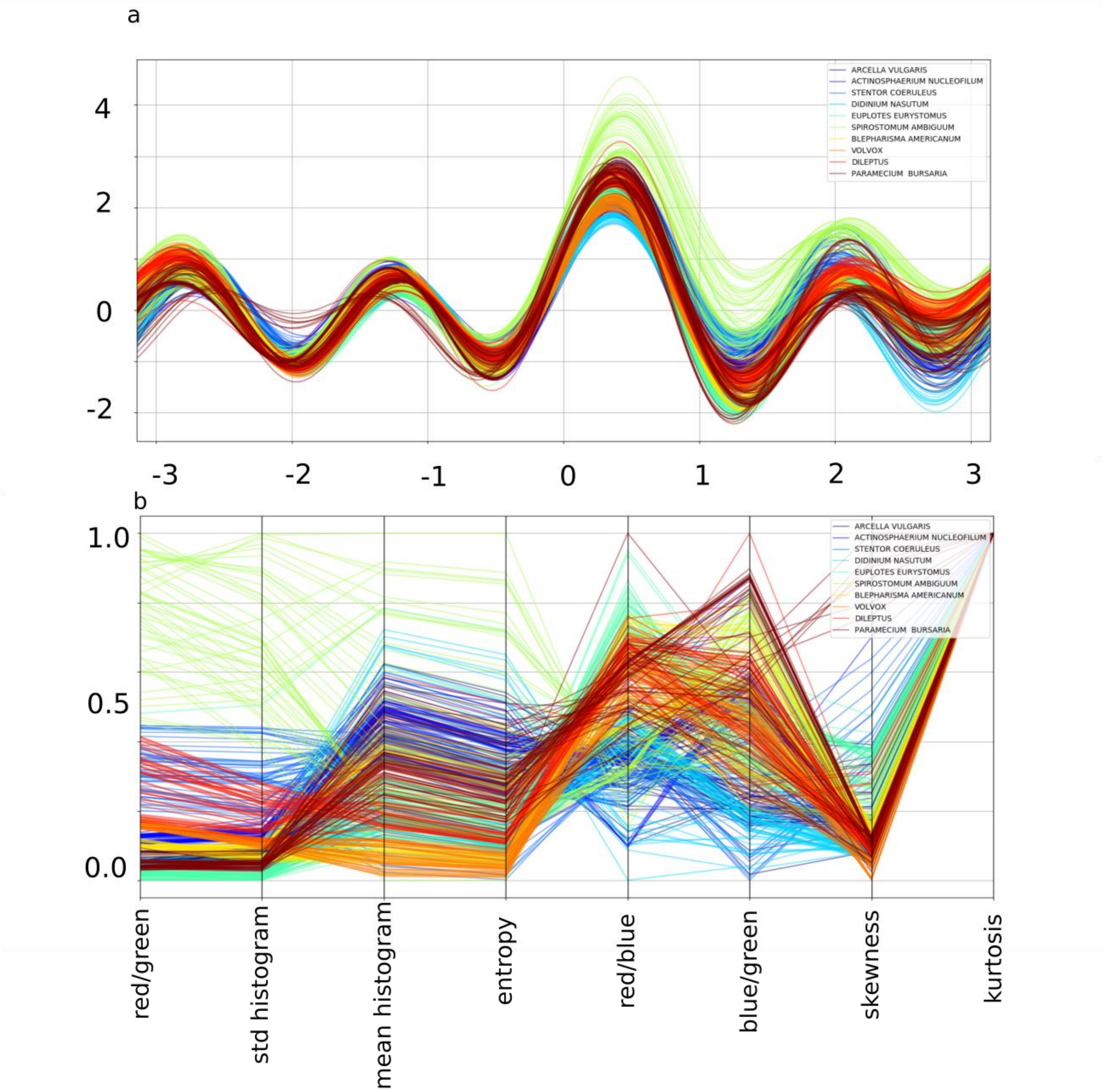


**Fig S9. Multi-dimensional representation for the features extracted from the gr ay values histogram. a** Andrew's curve**. b** Parallel coordinates

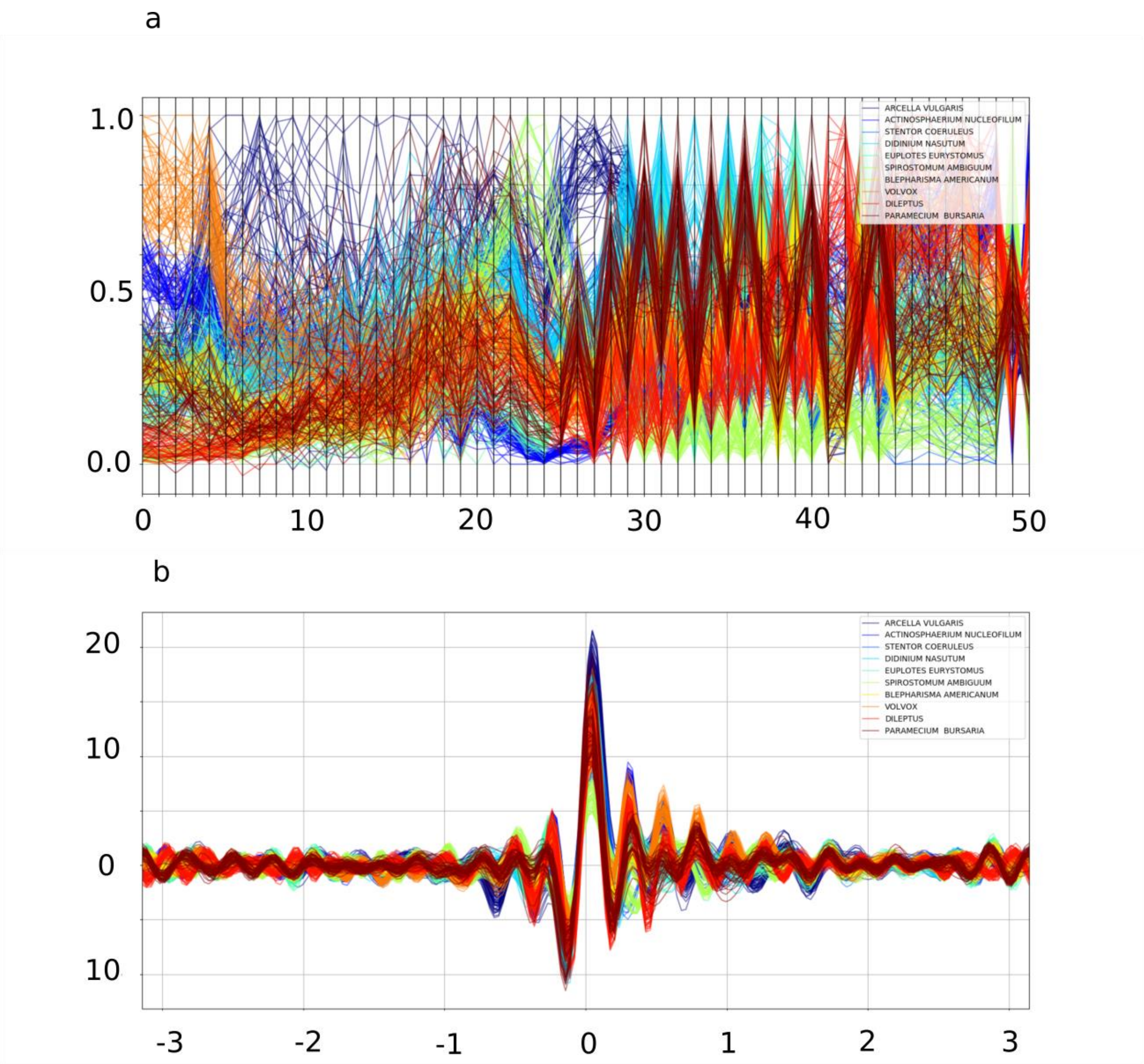


**Fig S10. Multi-dimensional representation for the LBP subset of features. a** Andrew’s curve**. b** Parallel coordinates

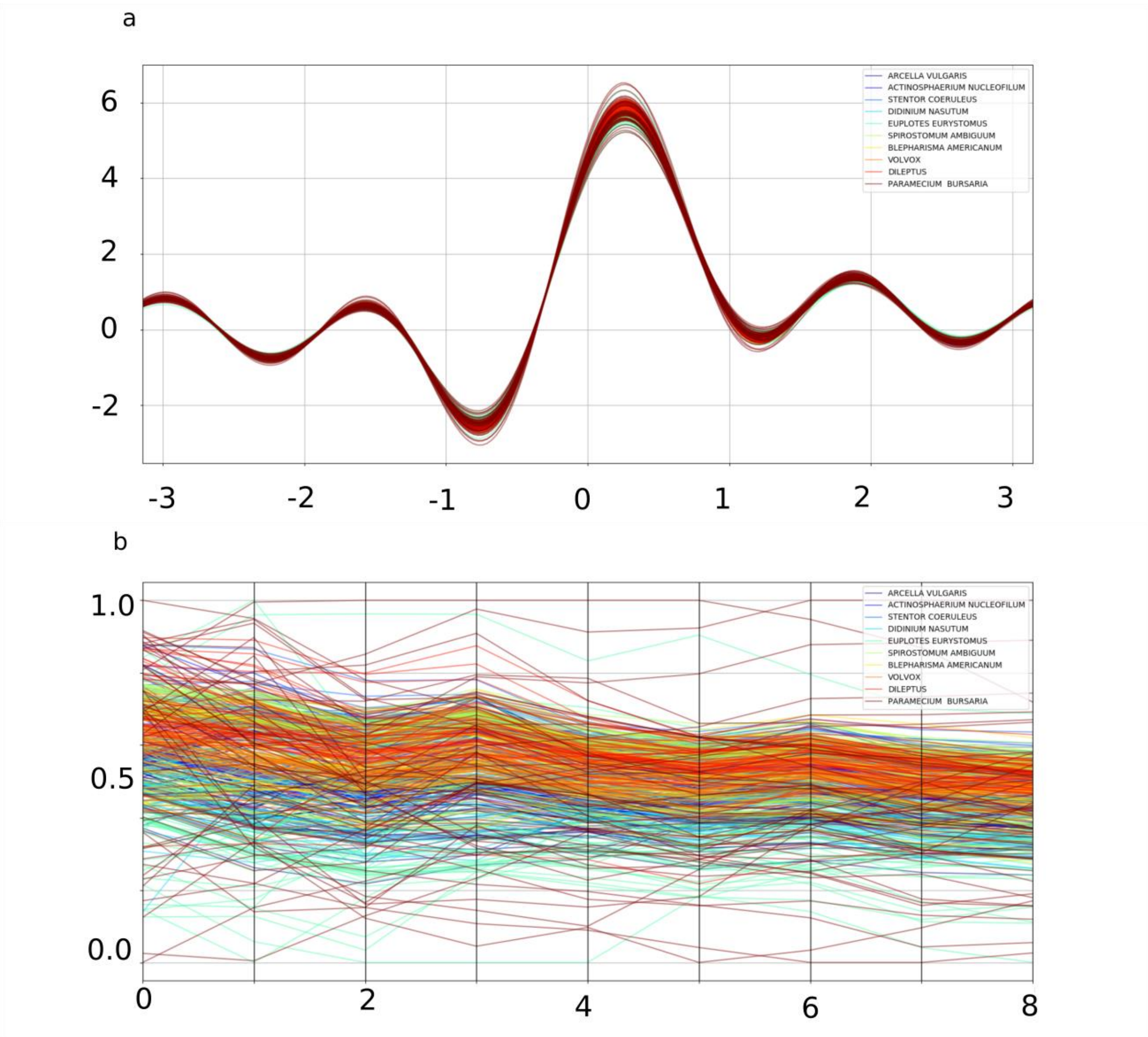


**Fig S11. Multi-dimensional representation for the Fourier Descriptors subset of features. a** Andrew's curve**. b** Parallel coordinates

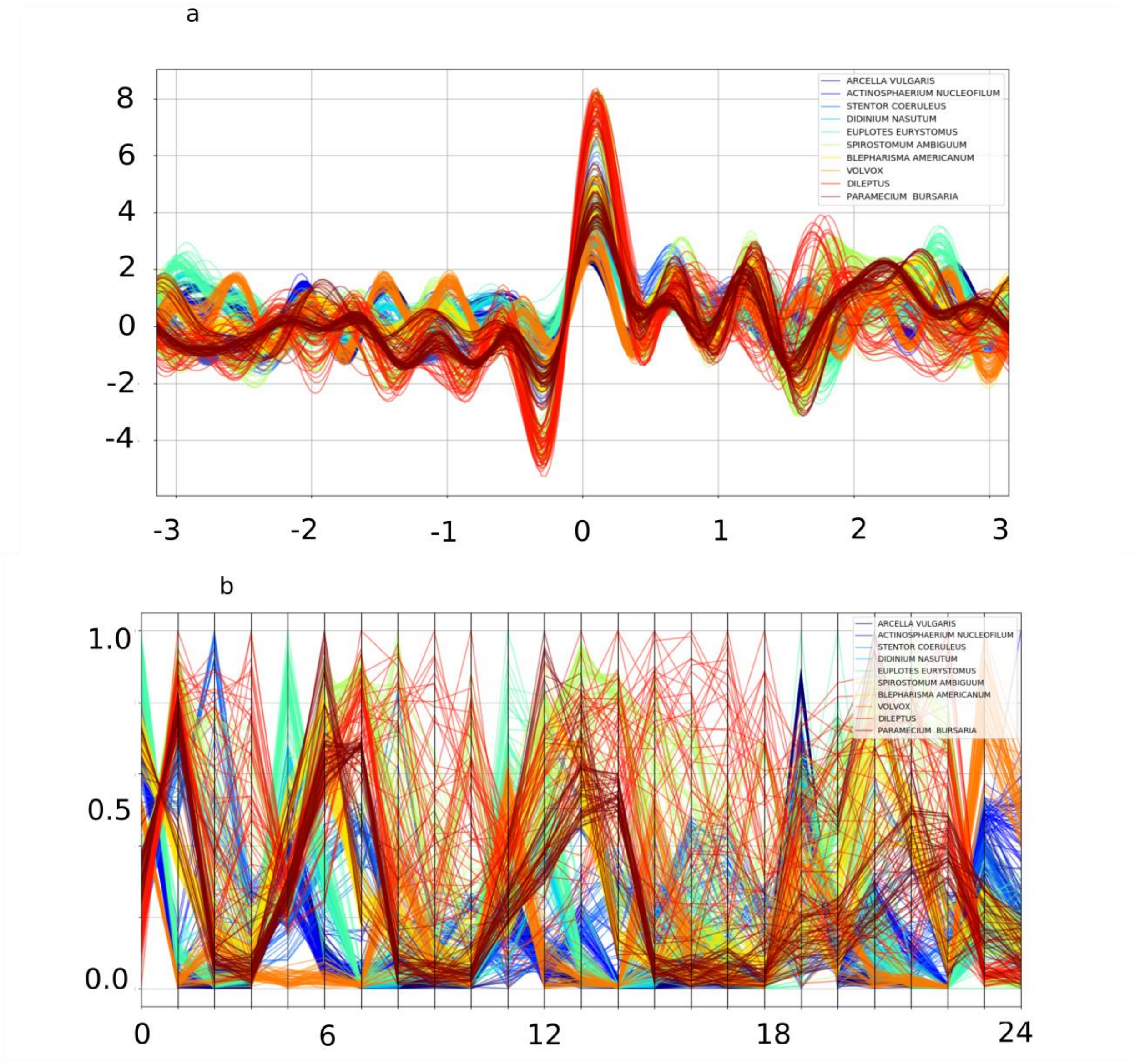


**Fig S12. Multi-dimensional representation for the Zernike moments subset of features. a** Andrew's curve**. b** Parallel coordinates

#### FEATURES RANKING

We used an algorithm based on decision trees to define a ranking for the 131 designed shape descriptors, based on their capacity to separate the 10 plankton classes used for training of the developed system. Fig S13 shows the results in form of a histogram. It is possible to appreciate that the geometric, the first two Hu-moments and the Haralick descriptors are the most important features in separating the training species. On the other hand, the Fourier descriptors seem to be the less interesting features for our specific problem. However, removing them caused a little decrease in the total accuracy, thus, we included them in the final set of designed descriptors.

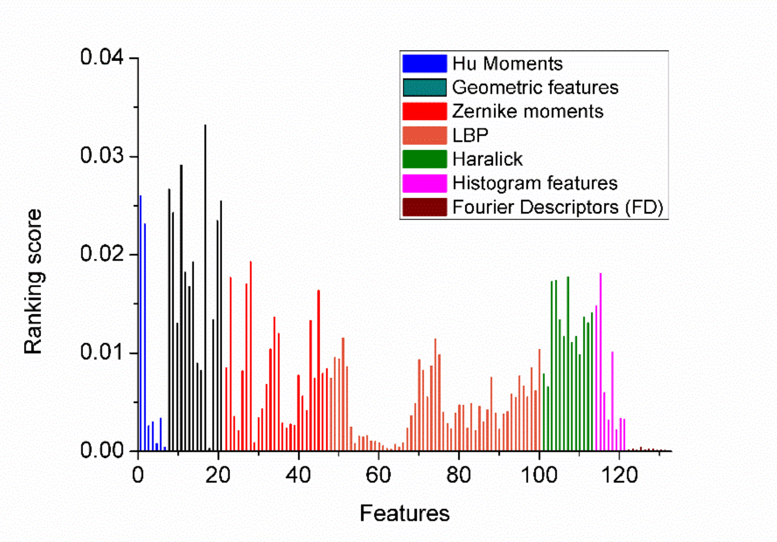


**Fig S13 Histogram reporting the normalized ranking score for the set of designed descriptors.**

#### FROM OBSERVATION TO IDENTIFICATION

In this paragraph, we will describe briefly how the DEC detectors are assembled together into the architecture designed to provide a continuous and real-time environmental monitoring (see fig S14). Specifically, each observation (i.e., the array of features extracted from the detected plankton cell image into the acquired videos) is given as input to each of the N detectors correspondent to the N species used for the training. A sample is assigned to class *i*, if and only if the *i* class detectors output is 1 and all the other detectors anomaly output is equal to 1. A sample is classified as anomaly in case that the anomaly output for all the implemented detectors are equal to 1. Finally, if more than one detector recognizes the observation as belonging to the correspondent class (i.e., if more than one detector's class output is equal to 1), the observation is assigned to the most activated detector, and labeled as retrain sample. These retrain samples will be used to retrain the correspondent detectors to reach a higher accuracy.

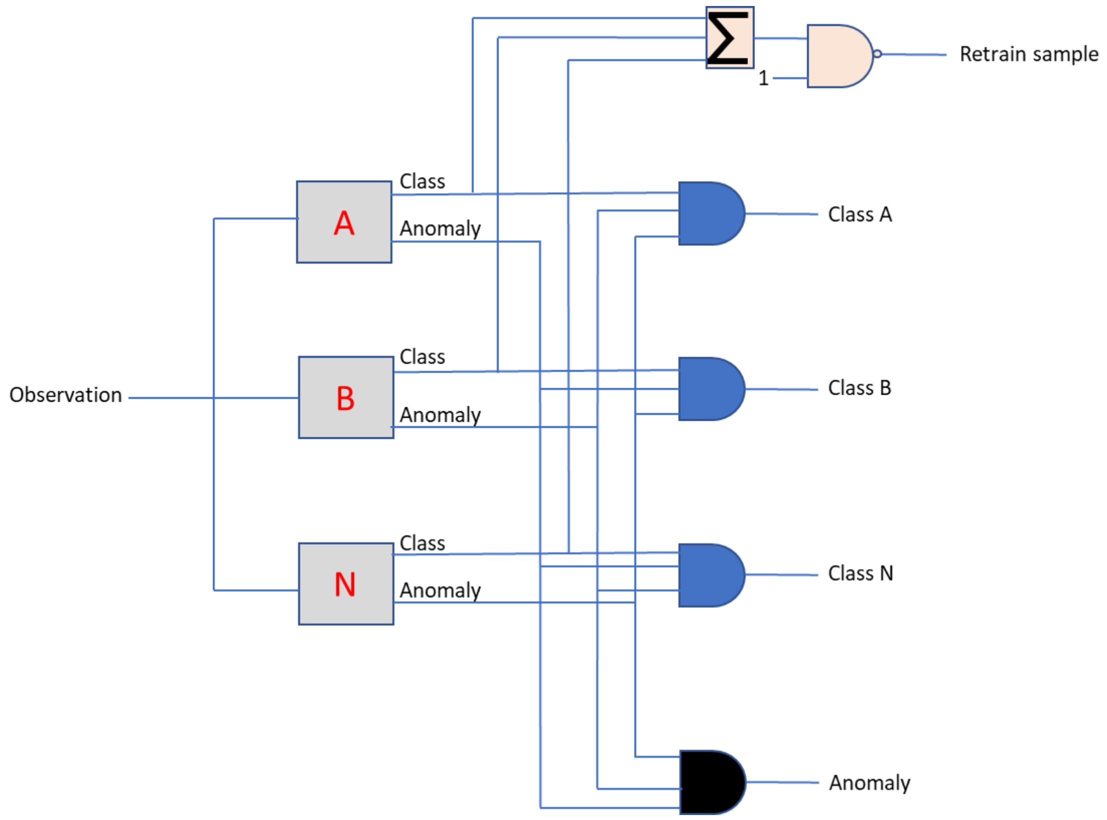


**Fig S14. Schematic work flow describing how an observation is associated to the three possible outpus of the developed system: retraining class, anomaly or belonging to a trained class**

### Pipeline Performance

#### COMPUTATIONAL TIME ON RASPERRY-PI

We performed a test on a raspberry-pi 3.0 to determine the computational time necessary to classify a sample plankton image from the acquired video. Table S1 summarizes the results, divided by computational module. In detail, we computed the requested time to detect and crop the image, to extract the array of features and to test the resulting features vector by means of the DEC detector. Here we report the results relative to the analysis of 1 seconds of videos (30 frames) containing two plankton cells (*Stentor coeruleous*). The total average computational time for one sample is 0.64 seconds. The computational time increases of about 0.05 seconds for each DEC detector included into the analysis.

| Pipeline Module | Computational time (s) |
|---|---|
| VIDEO ACQUISITION/DETECTOR | 0.15 ± 0.02 |
| FEATURE EXTRACTOR | 0.45 ± 0.04 |
| DEC SAMPLE TESTING | 0.04 ± 0.01 |

**Table S1. Computational time on raspberry pi for the analysis of one sample.** The standard deviation is computed among the objects contained into the 60 frames of the analyzed video.

#### COMPUTATIONAL TIME ON PC

We used a PC with CPU quad core i7 2.9 GhZ, 32 GB of RAM and GPU NVIDIA GeForce 940 MX with 2 GB of RAM. The computational time requested for the image processor module applied to the 40 species included into the WHOI dataset is around 49 seconds. The total computational time requested for the image processor module applied to the 100 images for each of the 40 species included into the WHOI dataset is around 49 seconds. The features extraction module for the same dataset requires 720 seconds (around 0.18 seconds per sample). The partitioning algorithm requires 32 seconds for the complete dataset, while the classification module based on random forest required  31 seconds (0.007 seconds per sample).

##### TIME REQUIRED FOR DEC-DETECTORS TRAINING ON PC

We trained the DEC detectors using the 500 training images for each of the ten species included into the lensless microscope dataset. The average time for training (see main text for training process details) is $t = 39.2 \pm 3.4$ seconds.

#### References


1. Bezdek, J. C., Ehrlich, R. & Full, W. FCM: The fuzzy c-means clustering algorithm. *Comput. Geosci.* **10**, 191–203 (1984).
2. Xie, X. L. & Beni, G. A Validity Measure for Fuzzy Clustering. *IEEE Trans Pattern Anal Mach Intell* **13**, 841–847 (1991).
3. MacQueen, J.B. (1967) Some Methods for Classification and Analysis of Multivariate Observations. Proceedings of 5-th Berkeley Symposium on Mathematical Statistics and Probability, 1, 281-297. - Open Access Library. Available at: http://www.oalib.com/references/14746902. (Accessed: 11th November 2018)
4. Lloyd, S. Least squares quantization in PCM. *IEEE Trans. Inf. Theory* **28**, 129–137 (1982).
5. Dunn, J. C. A Fuzzy Relative of the ISODATA Process and Its Use in Detecting Compact Well-Separated Clusters. *J. Cybern.* **3**, 32–57 (1973).

6. Pattern Recognition with Fuzzy Objective Function Algorithms | James C. Bezdek | Springer. Available at: https://www.springer.com/us/book/9781475704525. (Accessed: 11th November 2018)
7. Reynolds, D. A. Gaussian Mixture Models. in *Encyclopedia of Biometrics* (2009). doi:10.1007/978-0-387-73003-5_196
8. Huang, Z. & Leng, J. Analysis of Hu's moment invariants on image scaling and rotation. *2010 2nd Int. Conf. Comput. Eng. Technol.* **7**, V7-476-V7-480 (2010).
9. Yang, Z. & Fang, T. On the Accuracy of Image Normalization by Zernike Moments. *Image Vis. Comput* **28**, 403–413 (2010).
10. Coelho LP. Mahotas. Open source software for scriptable computer vision. *Journal of Open Research Software.* (2013). doi:DOI: http://doi.org/10.5334/jors.ac
11. Haralick, R. M., Shanmugam, K. & Dinstein, I. Textural Features for Image Classification. *IEEE Trans. Syst. Man Cybern.* **SMC-3**, 610–621 (1973).
12. Ojala, T., Pietikainen, M. & Maenpaa, T. Multiresolution gray-scale and rotation invariant texture classification with local binary patterns. *IEEE Trans. Pattern Anal. Mach. Intell.* **24**, 971–987 (2002).
13. Huang, D., Shan, C., Ardabilian, M., Wang, Y. & Chen, L. Local Binary Patterns and Its Application to Facial Image Analysis: A Survey. *IEEE Trans. Syst. Man Cybern. Part C Appl. Rev.* **41**, 765–781 (2011).
14. Osowski, S. & Nghia, D. D. Fourier and wavelet descriptors for shape recognition using neural networks—a comparative study. *Pattern Recognit.* **35**, 1949–1957 (2002).
15. Zhang, D. & Lu, G. A Comparative Study on Shape Retrieval Using Fourier Descriptors with Different Shape Signatures. in (2001).

16. Inselberg, A. The plane with parallel coordinates. *Vis. Comput.* **1**, 69–91 (1985).

17. Andrews, D. F. Plots of High-Dimensional Data. *Biometrics* **28**, 125–136 (1972).

18. Ho, T. K. Random decision forests. in *Document analysis and recognition, 1995., proceedings of the third international conference on* **1**, 278–282 (IEEE, 1995).

19. Genuer, R., Poggi, J.-M. & Tuleau, C. Random Forests: some methodological insights. *ArXiv08113619 Stat* (2008).

20. Breiman, L. Random Forests. *Mach. Learn.* **45**, 5–32 (2001).

21. Random forest algorithm for classification of multiwavelength data - IOPscience. Available at: http://iopscience.iop.org/article/10.1088/1674-4527/9/2/011. (Accessed: 11th November 2018)

22. Haykin, S. *Neural Networks: A Comprehensive Foundation*. (Prentice Hall PTR, 1994).

23. Schölkopf, B., Platt, J. C., Shawe-Taylor, J., Smola, A. J. & Williamson, R. C. Estimating the support of a high-dimensional distribution. *Neural Comput.* **13**, 1443–1471 (2001).
